# Old mice fail to integrate a memory update into an existing hippocampal engram

**DOI:** 10.64898/2026.06.22.733721

**Authors:** Chad A. Brunswick, Annie G. Defina, Trinity A. Wood, Derek J. Baldwin, Alexandria R. McKenna, Avery R. Sicher, Cyrus R. Marwaha, Dakota F. Brockway, Shoko Murakami, Gretchen C. Pifer, Chad W. Smies, Nicole A. Crowley, Janine L. Kwapis

## Abstract

Existing memories can be updated by the presentation of new information during memory retrieval. Memory updating is impaired with age, and recent reports indicate this process is more susceptible to age-related impairments than the formation of new memories. However, the neuronal mechanisms underlying age-related updating impairments are unknown. Here, we investigated how memory engrams within the dorsal hippocampus encode a memory update in the young and old brain. We found that old mice tended to re-engage a smaller proportion of the original memory engram during the update session and chemogenetically increasing the activation of this engram alleviated age-related updating deficits. A range of therapeutically relevant behavioral and pharmacological approaches promoting re-engagement of the training engram also improved memory updating in old mice. Together, these results identify a novel mechanism by which memory updating is impaired with age and expand our understanding of how the brain organizes related information.

## Introduction

Memory systems allow prior experience to shape future behavior. As part of this process, individual memories must sometimes be updated to reflect the most recent and relevant contingencies.^1,2^ Although memory updating is thought to occur through the molecular process of reconsolidation,^3–6^ little is known about how neural representations change during a memory update. Understanding the mechanisms of memory updating is important, as this process is impaired by aging, with age-related updating deficits reported in both humans^7–10^ and rodents.^11,12^ Notably, updating deficits seem to occur earlier and more readily with age than deficits in new memory formation,^12–14^ indicating this process might be particularly susceptible to the effects of aging. Nonetheless, little research to date has explored the underlying mechanisms of age-related updating impairments, and the field has overwhelmingly focused on understanding *de novo* memory formation.

There is substantial evidence that memories are sparsely encoded by distinct neuronal ensembles—often referred to as engrams—that are highly active during both initial learning and subsequent memory recall.^15–19^ Further support for the engram hypothesis stems from work demonstrating that artificial activation of engram neurons is sufficient to induce memory recall,^20,21^ while inhibiting or ablating these cells disrupts memory recall.^19,22^ Selective activation of context-encoding engram neurons within the dorsal hippocampus (DH) during exposure to a fearful stimulus is sufficient to update this contextual memory to include a fearful component, indicating an important role for neuronal engrams in memory updating.^23^

Recent work suggests that a fundamental mechanism by which two related memories are linked to one another is through engram co-allocation, wherein these memories are represented by overlapping ensembles of neurons.^24–27^ However, studies of engram co-allocation thus far have exclusively examined memories that are related to one another by virtue of occurring close together in time. It reasons that a similar co-allocation mechanism would underlie the process of memory updating, such that the engram encoding memory A would be co-allocated with the engram encoding the updated memory A’. Indeed, this mechanism has been suggested in the literature^15,28^ but not yet formally examined in a memory updating task. Here, we use our recently described Objects in Updated Locations (OUL; see Methods)^11,29^ memory updating task to investigate the engram dynamics underlying memory updating and how these dynamics are affected by aging.

## Results

### dCA1 engram co-allocation during updating is disrupted in the aging brain

First, we used a viral TetTag approach^23,30^ to track memory engram dynamics in dorsal CA1 (dCA1) during memory updating. We chose to look at dCA1 as this region is critical for spatial memory,^31^ necessary for updating in OUL,^11^ and a known site of engram co-allocation during memory linking.^24,26^ To investigate memory updating, we used a variant of the OUL task consisting of six days of habituation followed by three days of training, an update day, and a test day.^11,32^ The initial three days of training are necessary to overcome age-related impairments in memory formation and allow us to compare updating performance between young and old mice without the confounding effects of differences in initial memory formation.

Young adult and old mice were bilaterally infused with the TetTag viral cocktail while kept on Dox, and two weeks later underwent the OUL protocol (**Fig. 1A**). To label the initial training engram, mice were taken off Dox 48 hrs prior to the first OUL training session and returned to the Dox diet immediately after this session to close the tagging window. One hour after the OUL update session, animals were sacrificed and the update engram was labeled with immunohistochemistry for c-Fos (**Fig. 1B**). No difference was detected between young adult or old mice in the total number of dCA1 neurons (**Fig. 1C**), the relative size of the training engram (**Fig. 1D**), or the relative size of the update engram (**Fig. 1E**). However, we found that young adult mice re-engaged a significantly higher proportion of the initial dCA1 training engram during the update session relative to old mice (**Fig. 1F**), exhibited a higher Similarity Index^33,34^ between these two engrams (**Fig. 1G**), and also had more double positive c-Fos^+^GFP^+^ cells overall (**Extended Data Fig. 2C**). Therefore, old mice show reduced co-allocation between the update and the original memories compared to young mice, which may explain their failure to properly learn the update.

**Figure 1.**
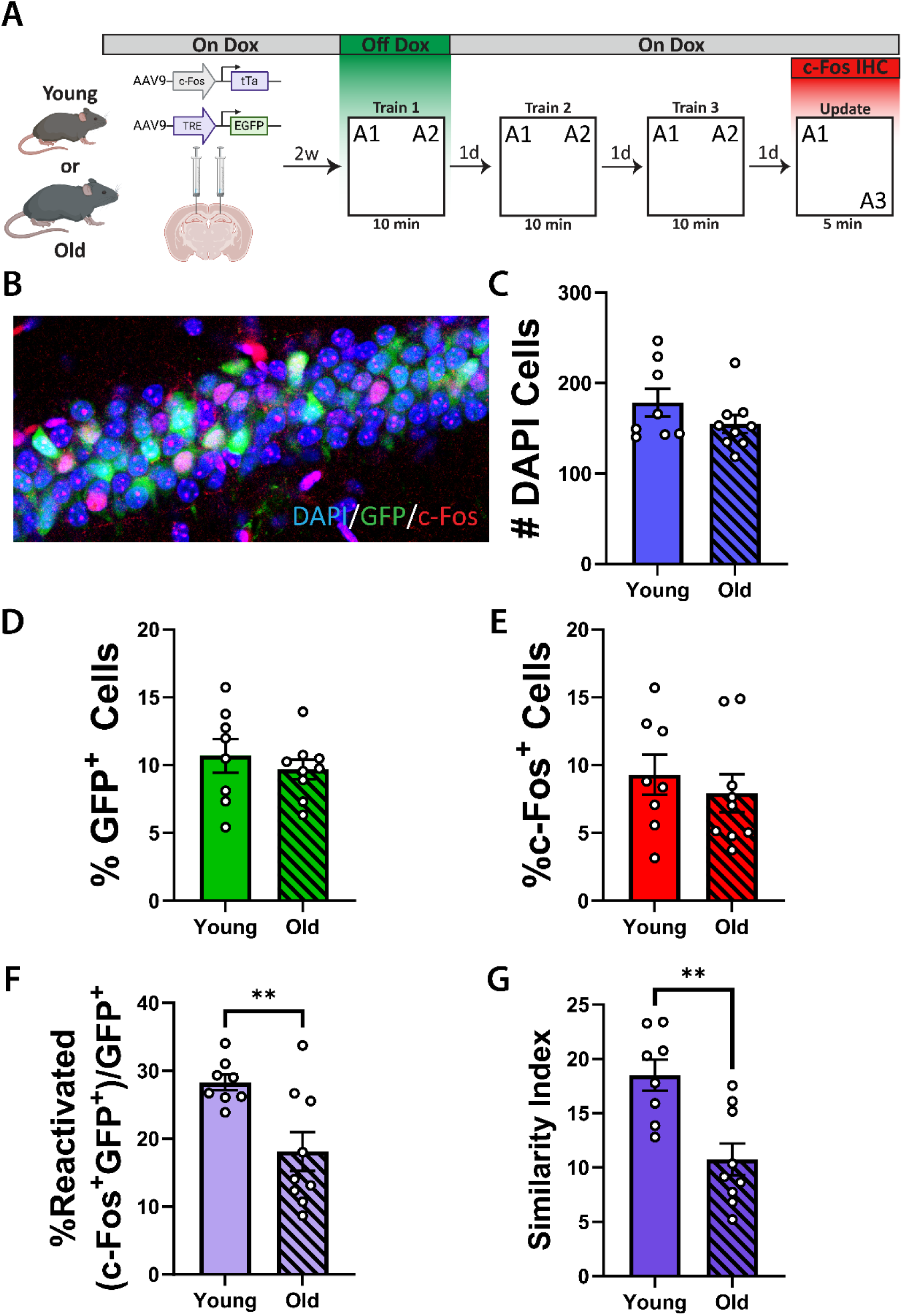
Hippocampal engram co-allocation during updating is decreased in old mice. **A.** Experimental schematic. Animals were injected with the dual virus TetTag system two weeks prior to engram tagging. Mice were removed from Dox 48 hrs prior to the first day of training to label the training engram with GFP and were then promptly restored to Dox. **B.** Representative image of GFP (green) and c-Fos (red) staining in dCA1. **C.** Similar numbers of total cells were counted per stack for each group (Mann-Whitney test: U = 26, p = 0.37). **D.** Percentage of GFP^+^ cells did not differ between age groups (unpaired Student’s t-test: t_15_ = 0.72, p = 0.48). **E.** Percentage of c-Fos^+^ cells did not differ between age groups (unpaired Student’s t-test: t_15_ = 0.66, p = 0.52). **F**. Reactivation rate of the training engram during updating was significantly higher in young mice compared to old mice (Welch’s t-test: t_10.45_ = 3.30, p = 0.0075). **G.** Similarity Indices comparing the training and update engrams were significantly higher in young mice compared to old mice (unpaired Student’s t-test: t_15_ = 3.74, p = 0.0020). All data are presented as mean ± SEM. n = 8 (4F), 9 (5F).

In both young adult and old mice, we observed intact preference for the A3 position during updating (**Extended Data Fig. 2A**) and engram re-engagement above chance levels (**Extended Data Fig. 2C**), together indicating that both groups of animals successfully retrieved the training memory during updating. Additionally, when we tested memory for the update just 75 minutes after updating, during a transcription-independent short-term memory window, old mice showed no deficits in their recollection of the update session (**Extended Data Fig. 3**). These results demonstrate that old mice show appropriate motivation and novelty detection during updating and acquire the updated information normally but fail to properly reconsolidate this information into long-term memory observable the following day.

### Activating the dCA1 training engram improves memory updating in old mice

Next, we investigated whether we could alleviate age-related deficits by specifically targeting this deficit in co-allocation. We used two distinct viral approaches to bias the recruitment of training engram neurons into the update engram. As excitable neurons are preferentially allocated to newly formed memory engrams,^35^ we expressed the excitatory designer receptor exclusively activated by designer drugs (DREADD) hM3Dq^36^ selectively in dCA1 training engram cells and then activated these neurons during updating to bias the recruitment of these neurons into the update engram, thereby increasing co-allocation.

First, we used an allocate-and-manipulate approach^18,37^ to increase co-allocation by biasing specific dCA1 neurons into both the training and update engrams. Old mice were administered a low titer of either AAV9-hSyn-hM3Dq-mCherry or AAV9-hSyn-mCherry to dCA1 two weeks prior to undergoing the OUL protocol (**Fig. 2A**). This low titer (5.0 × 10^10^ GC/mL) was chosen to transduce only around 10% of neurons in dCA1 (**Fig. 2B**), approximating the proportion of neurons allocated to a memory engram (e.g., in **Fig. 1D**).^22,24^ To activate hM3Dq and bias transduced neurons into the training engram, mice received i.p. injections of clozapine-n-oxide (CNO) 30 min prior to each training session to repeatedly activate this population of cells. Mice received an identical injection of CNO 30 min prior to the OUL update, thereby biasing these training engram neurons into the update engram as well. On the test day, mice received a saline injection 30 min prior to being placed into the OUL arena. To control for off-target effects of CNO,^38^ both groups of animals received identical CNO injections and differed only by the viral construct administered. The hM3Dq DREADD was validated through patch-clamp electrophysiology (**Extended Data Fig. 8A,B**).

**Figure 2.**
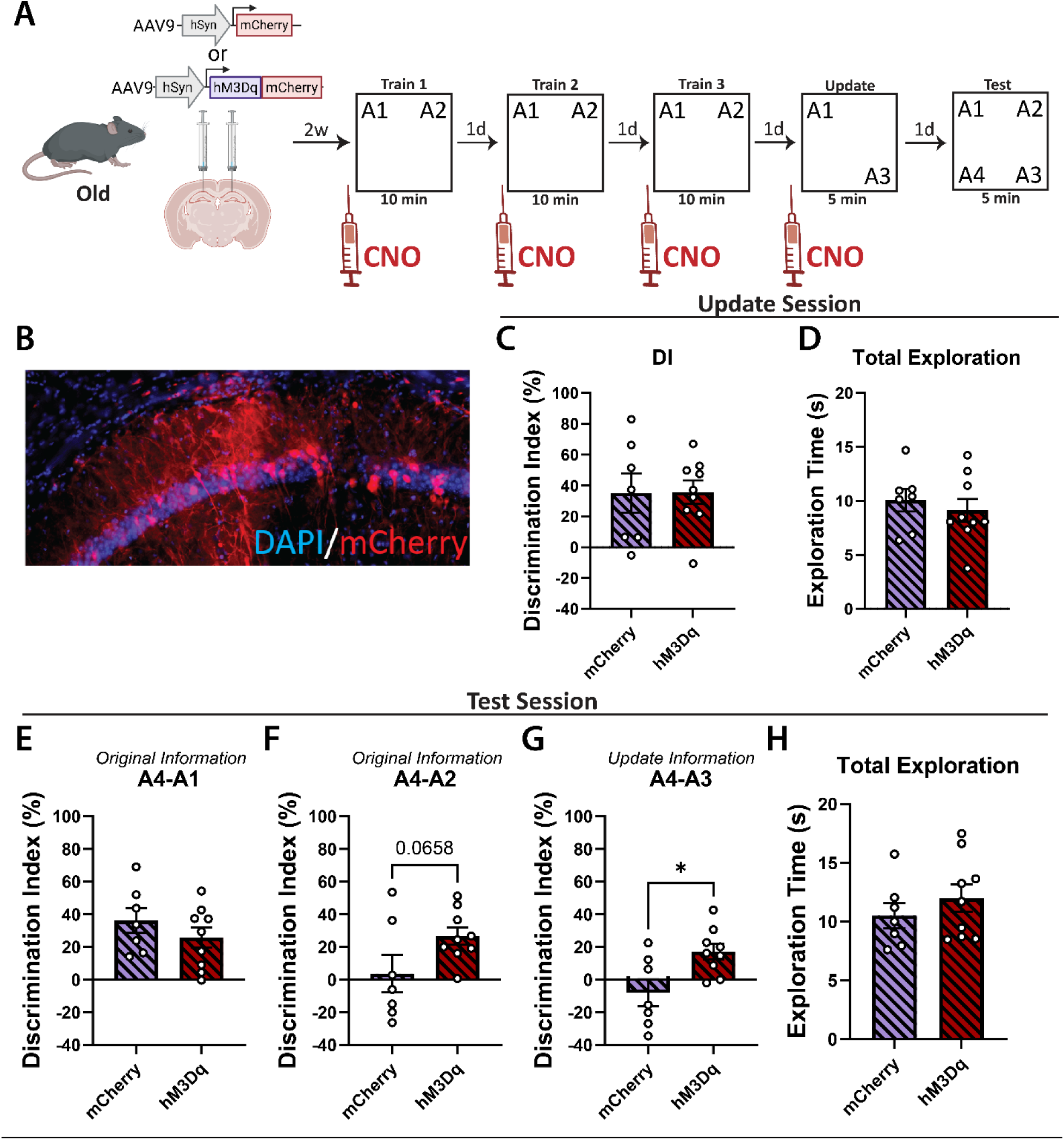
Repeated activation of the same dCA1 ensemble alleviates age-related updating impairments. **A.** Experimental schematic. Old mice were administered a low titer of either AAV9-hSyn-hM3Dq-mCherry or AAV9-hSyn-mCherry two weeks prior to OUL training. CNO was administered 30 min prior to each training session as well as the update session. **B.** Representative dCA1 injection, illustrating approximately 10% of dCA1 neurons expressing mCherry. **C.** Discrimination indices for training memory assessed during the update session. Both groups of animals showed intact memory for the initial training (one-sample t-tests of A3-A1 DIs against 0: mCherry: t_6_ = 2.76, p = 0.033; hM3Dq: t_8_ = 4.73, p = 0.0015), with no differences between groups (unpaired Student’s t-test: t_14_ = 0.041, p = 0.97). **D.** Total exploration time during the update session. No differences were observed between groups (t_14_ = 0.54, p = 0.62). **E.** During the test session, both groups of mice demonstrated intact memory for training position A1 (one-sample t-tests of A4-A1 DIs against 0: mCherry: t_6_ = 4.75, p = 0.032; hM3Dq: t_8_ = 4.07, p = 0.0036), with no differences between groups (unpaired Student’s t-test: t_14_ = 1.06, p = 0.30). **F.** Only the hM3Dq-expressing mice exhibited intact memory for training position A2 at test (one-sample t-tests of A4-A2 DIs against 0: mCherry: t_6_ = 0.31, p = 0.76; hM3Dq: t_8_ =5.12, p = 0.0009), though a direct comparison revealed only a trend towards a significant difference between the groups (unpaired Student’s t-test: t_14_ = 2.00, p = 0.066). **G.** At test, only mice in the hM3Dq group showed memory for the update position A3 (one-sample t-tests of A4-A3 DIs against 0: mCherry: t_6_ = 0.98, p = 0.36; hM3Dq: t_8_ =3.60, p = 0.0070), and these mice performed significantly better than mice in the mCherry group (unpaired Student’s t-test: t_14_ = 2.80, p = 0.014). **H.** These effects were not driven by any differences in total exploration during the test session (unpaired Student’s t-test: t_14_ = 0.92, p = 0.38). All data are presented as mean ± SEM. n = 7 (2F), 9 (4F).

Animals habituated to the environment as expected (**Extended Data Fig. 4A)** and demonstrated typical behavior during the three training sessions (**Extended Data Fig. 4B,C**). During the update session, both groups displayed intact memory for the training sessions (**Fig. 2C**) and usual typical exploration times (**Fig. 2D**). No difference in behavior was observed between the groups during the update session, indicating that activation of the training engram here fails to further improve memory for the training sessions. This is likely due to a ceiling effect, as three days of training typically produces robust memories during the update session, even for old mice.^11,32^

At test, we observed intact memory in all animals for the training location A1 (**Fig. 2E**). Interestingly, mCherry control mice failed to remember location A2 at test, while the hM3Dq animals successfully did so (**Fig. 2F**) indicating a decrease in the retroactive interference occasionally observed in this task.^11^ Finally, only hM3Dq-expression mice showed intact memory for the update location A3 at test (**Fig. 2G**), indicating that activation of the dCA1 training engram during the update session improves memory updating in old mice. These effects were not driven by changes in overall exploration time during the test session (**Fig. 2H**). Raw percent exploration time for each object during the test session is shown in **Extended Data Fig. 4D**.

To ask this question another way, we also employed a TetTag-DREADD approach to capture the engram endogenously engaged during training and selectively reactivate these cells during updating (**Fig. 3A**). Here, old mice were kept on Dox for one week prior to stereotaxic surgery where animals were administered AAV9-cFos-tTA alongside either AAV9-TRE-EGFP or AAV9-TRE-hM3Dq-EGFP in dCA1. Two weeks after viral surgery, animals underwent the OUL protocol. As before, animals were removed from Dox following habituation and left undisturbed for 48 hrs in their homecages until the first day of OUL training to selectively label the initial training engram (**Fig. 3B**). Immediately following the first training session, animals were restored to Dox and underwent the rest of the OUL protocol. On the update day, 30 min before the update session, animals received injections of CNO to selectively activate the tagged dCA1 training engram. Both groups of mice received CNO to minimize any confounding effects of drug administration, and the only difference between groups was whether animals received TetTag constructs containing hM3Dq-EGFP or EGFP alone (control animals).

**Figure 3.**
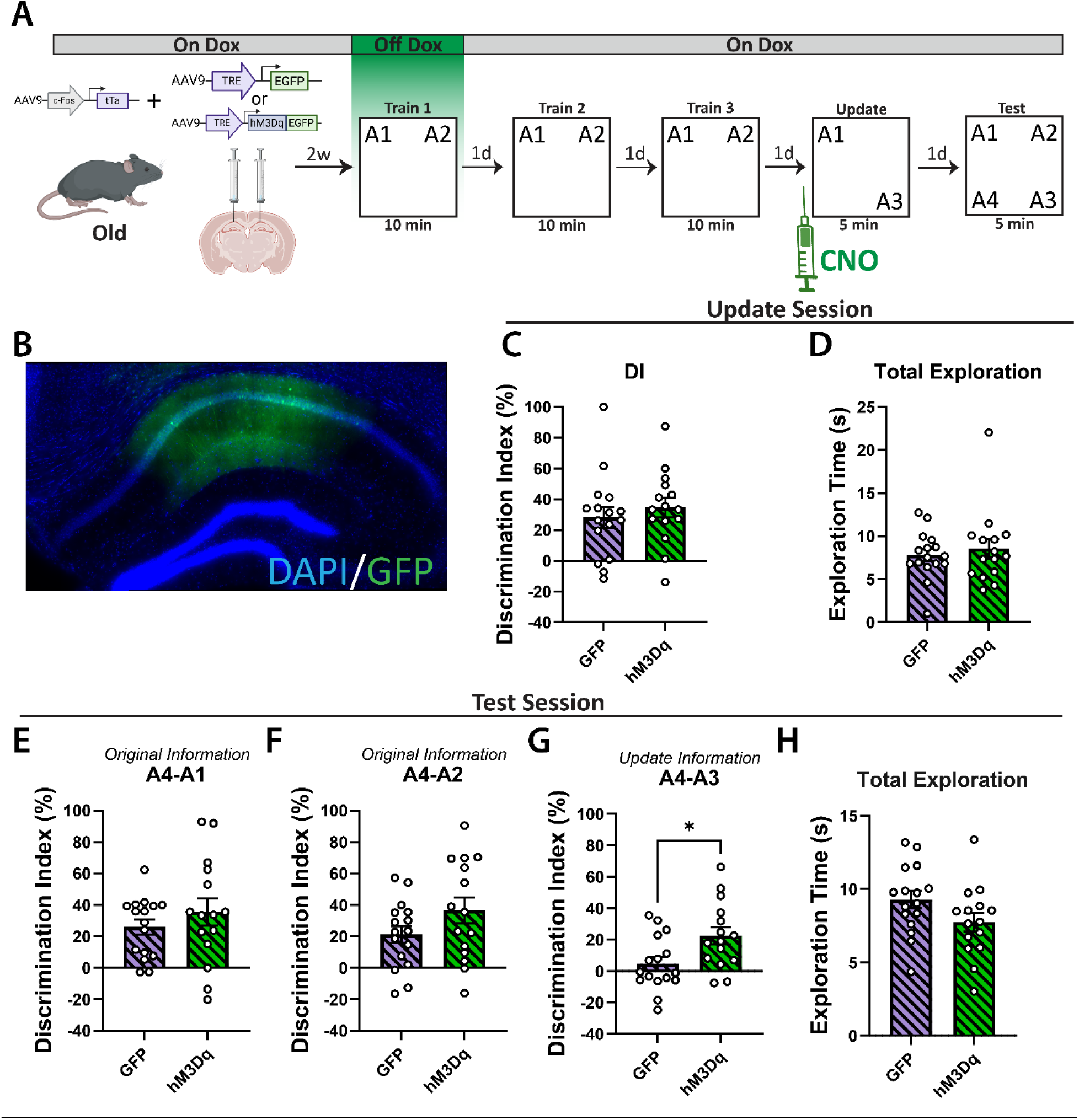
Activation of the training engram alleviates age-related updating impairments. **A.** Experimental schematic. Animals were injected with the dual virus TetTag system (expressing either hM3Dq-GFP or GFP alone) two weeks before engram tagging. Mice were removed from Dox 48 hrs prior to the first day of training to label the training engram with GFP and were then promptly restored to Dox. CNO was administered 30 min before the update session. **B.** Representative dCA1 injection. **C.** During the update session, both groups demonstrated a clear memory for training position A1 (one-sample t-tests of A3-A1 DIs against 0: GFP: t_15_ = 4.11, p = 0.0009; hM3Dq: t_14_ = 5.54, p < 0.0001), with no differences observed between groups (unpaired Student’s t-test: t_29_ = 0.67, p = 0.51). **D.** Both groups of mice also spent similar amounts of time exploring objects during the update session (unpaired Student’s t-test: t_29_ = 0.58, p = 0.57). **E.** At test, both groups of mice showed memory for position A1 from training (one-sample t-tests of A4-A1 DIs against 0: GFP: t_15_ = 5.47, p < 0.0001; hM3Dq: t_14_ = 4.07, p = 0.0011) with no differences between groups (Welch’s t-test: t_21.75_ = 0.96, p = 0.35). **F.** Both groups demonstrated intact memory for training position A2 at test (one-sample t-tests of A4-A2 DIs against 0: GFP: t_15_ = 3.94, p = 0.0013; hM3Dq: t_14_ = 4.46, p = 0.0005) with no differences between groups (unpaired Student’s t-test: t_29_ = 1.59, p = 0.12). **G.** Only mice in the hM3Dq group demonstrated memory for the update session (one-sample t-tests of A4-A3 DIs against 0: GFP: t_15_ = 1.03, p = 0.32; hM3Dq: t_14_ = 4.08, p = 0.0011), exhibiting significantly higher A4-A3 DIs than those in the GFP group (unpaired Student’s t-test: t_29_ = 2.60, p = 0.015). **H.** These effects were not driven by any differences in test session exploration (unpaired Student’s t-test: t_29_ = 1.79, p = 0.084), though hM3Dq mice trended towards slightly less exploration time overall. All data are presented as mean ± SEM. n = 16 (10F), 15 (8F).

All animals habituated to the environment as expected (**Extended Data Fig. 5A**). During the three training sessions, we observed similar behavior between both groups, with no differences in training DIs (**Extended Data Fig. 5B**) or exploration time (**Extended Data Fig. 5C**), and at the update session, both groups exhibited intact memory for the initial training (**Fig. 3C**). We observed no differences during updating between hM3Dq-expressing mice and EGFP-only controls in either memory for training or total exploration times (**Fig. 3C,D**), again indicating that three training sessions produce a robust memory in old mice.

During the test session, both groups exhibited intact memory for training location A1 (**Fig. 3E**). Both groups of old mice exhibited intact memory for training location A2 as well (**Fig. 3F**) with no significant differences detected between hM3Dq and control animals. Control EGFP-expressing mice failed to remember the update location A3 at test, while hM3Dq-expressing mice exhibited intact memory for this location (**Fig. 3G**), indicating that this procedure improved memory updating in these animals. As in the allocate-and-manipulate experiment, these results were not due to changes in total exploration time at test (**Fig. 3H**). Notably, hM3Dq mice were trending lower than EGFP controls, indicating a pattern of highly specific, focused exploration that would be expected to accompany robust memory retrieval. Raw percent exploration time for each object during the test session is shown in **Extended Data Fig. 5D**. Together, these experiments demonstrate that artificially encouraging neuronal co-allocation in the dorsal hippocampus ameliorates age-related impairments in spatial memory updating.

### Inhibiting the dCA1 training engram does not affect memory updating in young mice

Having demonstrated that activating the dCA1 training engram during updating is sufficient to improve memory updating in old mice, we next tried inhibiting the dCA1 training engram in young mice during updating to see if this would block memory updating. Here, we used a similar TetTag approach, now with the inhibitory receptor hM4Di^36^ downstream of the TRE promoter (**Fig. 4A**).

**Figure 4.**
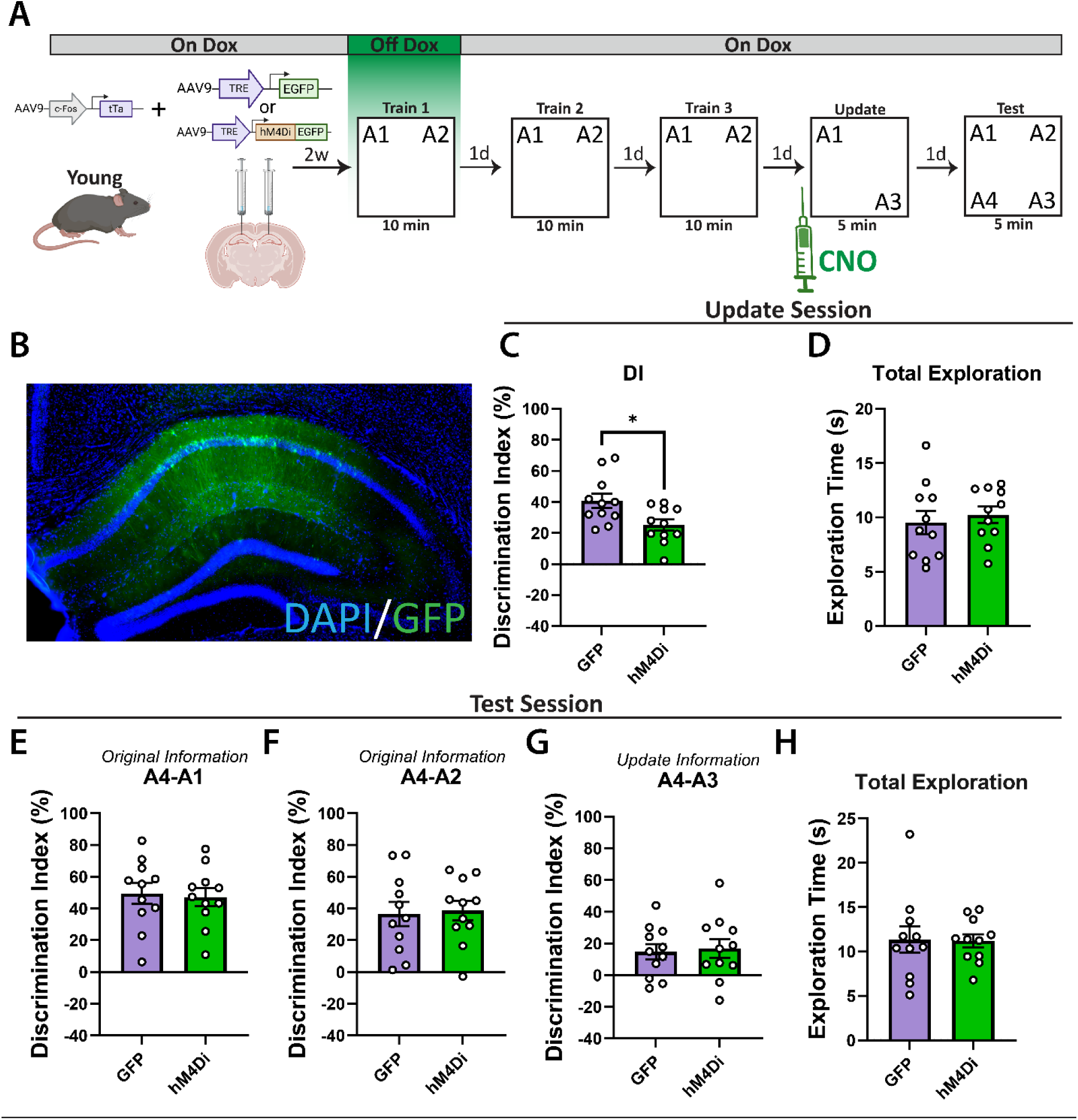
Inhibition of the training engram does not interfere with updating in young mice. **A.** Experimental schematic. Animals were injected with the dual virus TetTag system (expressing either hM4Di-GFP or GFP alone) two weeks prior to engram tagging. Mice were removed from Dox 48 hrs prior to the first day of training to label the training engram with GFP and were then promptly restored to Dox. CNO was administered 30 min prior to the update session. **B.** Representative dCA1 injection. **C.** During the update session, both groups of mice demonstrated intact memory for the initial training (one-sample t-tests of A3-A1 DIs against 0: GFP: t_10_ = 8.81, p < 0.0001; hM4Di: t_10_ = 7.18, p < 0.0001), though mice in the hM4Di group performed significantly worse than the GFP group, consistent with inactivation of the training engram (unpaired Student’s t-test: t_20_ = 2.68, p = 0.015). **D.** No differences were observed in object investigation during the update session (unpaired Student’s t-test: t_20_ = 0.54, p = 0.59). **E.** At test, both groups of mice demonstrated intact memory for training position A1 (one-sample t-tests of A4-A1 DIs against 0: GFP: t_10_ = 7.46, p < 0.0001; hM4Di: t_10_ = 8.31, p < 0.0001) with no differences observed between groups (unpaired Student’s t-test: t_20_ = 0.26, p = 0.80). **F.** Both groups also remembered position A2 from training (one-sample t-tests of A4-A2 DIs against 0: GFP: t_10_ = 4.80, p = 0.0007; hM4Di: t_10_ = 6.24, p < 0.0001), with no differences detected between groups (unpaired Student’s t-test: t_20_ = 0.22, p = 0.83). **G.** Both groups of mice also demonstrated intact memory for update position A3 (one-sample t-tests of A4-A3 DIs against 0: GFP: t_10_ = 2.99, p = 0.014; hM4Di: t_10_ = 2.75, p = 0.021), with no differences detected between groups (unpaired Student’s t-test: t_20_ = 0.27, p = 0.79). **H.** We additionally detected no differences in total investigation time at test (Mann-Whitney test: U = 56.5, p = 0.81). All data are presented as mean ± SEM. n = 11 (5F), 11 (5F).

Young adult mice were kept on Dox food for at least one week prior to being administered AAV9-cFos-tTa and either AAV9-TRE-hM4Di-EGFP or AAV9-TRE-EGFP (control animals) to dCA1. Following surgery, animals were maintained on Dox for two weeks and then underwent the OUL protocol. Mice were taken off Dox for the first training day to label the initial training engram (**Fig. 4B**), after which mice were placed back on Dox. On the update day, both groups of animals received intraperitoneal injections of CNO to activate the hM4Di receptor, selectively decreasing the excitability of just the dCA1 training engram. Both groups received CNO injections to minimize any confounding effects of drug administration. The hM4Di DREADD was validated through patch-clamp electrophysiology (**Extended Data Fig. 8C,D**).

Both groups of animals behaved similarly during habituation and training (**Extended Data Fig. 6A-C**). During the update session, (following CNO administration) we observed intact memory for the initial training in both the EGFP control mice and the hM4Di mice (**Fig. 4C**). However, hM4Di mice exhibited significantly lower A3-A1 DIs compared to EGFP controls, indicating that this approach successfully reduces the excitability of the dCA1 training engram and impairs memory retrieval during the update. We observed no differences in exploration time during the update (**Fig. 4D**).

At test, both groups of mice demonstrated intact memory for training locations A1 (**Fig. 4E**) and A2 (**Fig. 4F**). Notably, both groups of animals also demonstrated successful recall of update location A3 (**Fig. 4G**), indicating that this procedure failed to disrupt memory updating in young mice. We also observed no differences in total investigation time between both groups (**Fig. 4H**). Raw percent exploration time for each object during the test session is shown in **Extended Data Fig. 6D**.

We wanted to confirm that these null results were not simply a result of our experimental design. Although we saw decreases in memory recall during updating (**Fig. 4C**), and validated our DREADD (**Extended Data Fig. 8**), we wondered if the dose of CNO used might have been insufficient to fully inactivate these neurons, as some groups use a higher CNO dose for the hM4Di DREADD compared to the hM3Dq.^39^ Additionally, we reasoned that inactivating the ensemble from only one of the three training days might still permit co-allocation between the update engram and the rest of the training ensemble. Thus, we repeated this experiment with a higher dose of CNO (5 mg/kg), and only a single day of training (which we have previously shown young mice can learn without issue^11,29^), but we nonetheless observed similar results (**Extended Data Fig. 7**). We therefore were unable to find evidence that blocking co-allocation prevented successful memory updating in young mice.

### A behavioral intervention, reducing the interval between training and updating, improves updating only in old mice

Thus far, our results indicate that engram co-allocation is an important mechanism by which memory updates are reconsolidated into long-term storage and that this process is disrupted with aging. We wanted to leverage these results to identify potentially translatable strategies to improve memory updating in old age. First, we tested whether we could design a behavioral intervention that could improve memory updating in old age. To this end, we tested whether old mice could learn a memory update if we presented the update information shortly after the final training session. We reasoned that reducing the interval between training and updating would promote engram co-allocation, as this is a known mechanism of memory linking.^26,27,40^ Therefore, if memory updating is also dependent on engram co-allocation as our results suggest, then this procedure should bolster memory updating.

Old mice underwent the OUL paradigm as described except half of the mice were presented with the update session 75 min after the final training session (**Fig. 5A**). As usual, the test session was presented 24 hours following the update for all mice. We observed normal behavior during both habituation to the OUL arenas (**Extended Data Fig. 9A**) and the initial training sessions (**Extended Data Fig. 9B,C**). During the update session, both groups demonstrated intact memory for training (**Fig. 5B**), with no significant differences between groups. We observed a significant difference in investigation time during the update, with mice in the 75 min group investigating significantly more (**Fig. 5C**) than control mice, likely due to increased salience of the updated information at this short timepoint.

**Figure 5.**
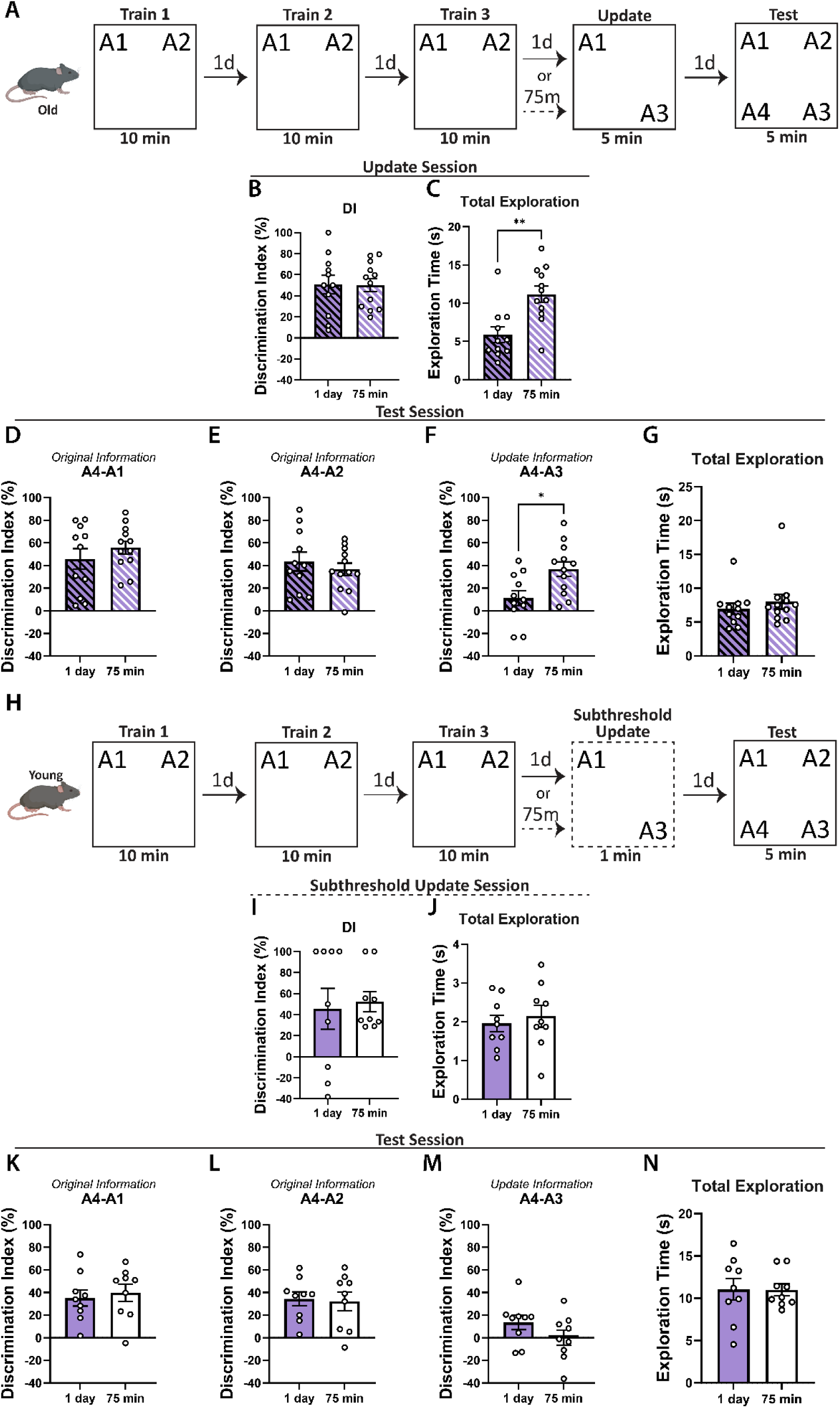
Presenting the update shortly after training improves updating in old mice but not in young mice. **A.** Old mice experimental schematic. Old mice underwent OUL and were given the update session 1 day or 75 min after the last training session. **B.** During the update session, both groups of old mice demonstrated intact memory for the initial training (one-sample t-tests of A3-A1 DIs against 0: 1 day: t_10_ = 5.58, p = 0.0002; 75 min: t_11_ = 8.01, p < 0.0001). No differences were observed between groups (unpaired Student’s t-test: t_21_ = 0.035, p = 0.97). **C.** Old mice in the 75 min group spent significantly more time investigating during the update session (unpaired Student’s t-test: t_21_ = 3.65, p = 0.0015). **D.** During test, both groups of old mice demonstrated intact memory for training position A1 (one-sample t-tests of A4-A1 DIs against 0: 1 day: t_10_ = 5.01, p = 0.0005; 75 min: t_11_ = 9.80, p < 0.0001), with no differences between groups (unpaired Student’s t-test: t_21_ = 0.96, p = 0.35). **E.** Both groups of old mice also remembered training position A2 (one-sample t-tests of A4-A2 DIs against 0: 1 day: t_10_ = 5.25, p = 0.0004; 75 min: t_11_ = 6.70, p < 0.0001), with no group differences (unpaired Student’s t-test: t_21_ = 0.69, p = 0.50). **F.** Only old mice in the 75min group remembered update position A3 (one-sample t-tests of A4-A3 DIs against 0: 1 day: t_10_ = 1.71, p = 0.12; 75 min: t_11_ = 5.64, p = 0.0002), performing significantly better than mice in the 1 day group (unpaired Student’s t-test: t_21_ = 2.75, p = 0.012). **G.** No differences occurred in test session exploration (unpaired Student’s t-test: t_21_ = 0.76, p = 0.46). **H.** Young mice experimental schematic. Young mice underwent OUL and were given a subthreshold update either 1 day or 75 min after the last training session. **I.** During the update session, both groups demonstrated intact memory for the initial training (one-sample t-tests of A3-A1 DIs against 0: 1 day: t_8_ = 2.35, p = 0.047; 75 min: t_8_ = 5.46, p = 0.0006), with no group differences (unpaired Student’s t-test: t_16_ = 0.31, p = 0.76). **J.** No differences were observed in update exploration time (unpaired Student’s t-test: t_16_ = 0.52, p = 0.61). **K.** At test, both groups demonstrated intact memory for training position A1 (one-sample t-tests of A4-A1 DIs against 0: 1 day: t_8_ = 4.91, p = 0.0012; 75 min: t_8_ = 5.26, p = 0.0008), with no differences between them (unpaired Student’s t-test: t_16_ = 0.45, p = 0.66). **L.** Both groups remembered training position A2 (one-sample t-tests of A4-A2 DIs against 0: 1 day: t_8_ = 5.52, p = 0.0006; 75 min: t_8_ = 3.87, p = 0.0047), with no group differences (unpaired Student’s t-test: t_16_ = 0.22, p = 0.83). **M.** Neither group remembered update position A3 (one-sample t-tests of A4-A3 DIs against 0: 1 day: t_8_ = 2.11, p = 0.068; 75 min: t_8_ = 0.0093, p = 0.99), with no differences between groups (unpaired Student’s t-test: t_16_ = 1.46, p = 0.16). **N.** No differences were observed in total exploration time at test (unpaired Student’s t-test: t_16_ = 0.0038, p = 0.97). All data are presented as mean ± SEM. n = 11, 12 (old); 9, 9 (young). All male.

At the retention test, all mice exhibited robust memory for locations A1 (**Fig. 5D**) and A2 (**Fig. 5E)** from the initial training. However, only mice in the 75 min group demonstrated intact memory for the update location A3 (**Fig. 5F**) indicating that presenting a memory update shortly after training is sufficient to ameliorate age-related memory updating impairments. This effect was not driven by changes in exploratory drive (**Fig. 5G**), and raw percent exploration time for each object during the test session is shown in **Extended Data Fig. 9D**.

We next investigated whether this shortened training-updating interval broadly improves memory updating or if this procedure uniquely alleviates age-related impairments. If this procedure enhances memory updating by restoring neuronal co-allocation as expected, this behavioral intervention should not improve updating in young mice, which already exhibit robust engram co-allocation (**Fig. 1**). Thus, we repeated this experiment with young adult mice given a subthreshold^32^ 1-min update session (**Fig. 5H**). In our young animals, we observed typical behavior during habituation (**Extended Data Fig. 9E**), and training (**Extended Data Fig. 9B,C**). During the subthreshold 1-min update session, mice had minimal time to explore the objects, and it is thus not possible to calculate a robust DI for this session. Nevertheless, both groups of mice demonstrated similar DIs (**Fig. 5I**) and total exploration times (**Fig. 5J**).

At test, both groups had intact memory for locations A1 (**Fig. 5K**) and A2 from training (**Fig. 5L**). However, neither group demonstrated intact memory for the update location A3, and no differences in memory for A3 were observed between groups (**Fig. 5M**), though total exploration times during test remained consistent for both groups of mice (**Fig. 5N**). Raw percent exploration time for each object during the test session is shown in **Extended Data Fig. 9H**. Thus, this temporal proximity strategy seems to specifically alleviate age-related deficits in memory reconsolidation and is not sufficient to transform a subthreshold update into long-term memory in young mice.

### An FDA-approved pharmacological intervention, inhibiting CCR5 with maraviroc, improves memory updating in old mice

We have established that engram co-allocation, a known mediator of memory linking, also mediates reconsolidation-driven memory updating, and our previous experiments suggest that leveraging mechanisms of memory linking (i.e., temporal proximity) might also enhance memory reconsolidation in old mice. Previous work on memory linking has demonstrated that this process is controlled in part by *Ccr5* (C-C chemokine receptor type 5) and its ligand *Ccl5* (Chemokine [C-C motif] ligand 5), both of which are negative regulators of memory co-allocation and expressed at increased concentrations in the aging DH.^26^ We therefore decided to explore if these genes also play a role in memory updating, as they have zet to be examined during this process. Elucidating the role of these genes in memory updating is of interest as CCR5 is a well-studied receptor with many FDA-approved antagonists,^41^ and leveraging this body of knowledge in the context of memory updating has important therapeutic implications for the treatment of cognitive decline.

First, we wanted to examine if there were any age-specific changes in hippocampal *Ccr5* or *Ccl5* expression following memory updating. To do so, we trained young adult and old mice in the OUL task as previously described and, following the last training session mice, were randomly assigned to one of three groups (**Fig. 6A**): homecage (HC), no update (NU), or update (U). Mice in the HC group were left in their homecage on the update day, mice in the NU group underwent a 5-min retrieval session identical to training (objects in locations A1 and A2) to control for retrieval-specific changes in gene expression, and mice in the U group underwent the typical 5-min OUL update session (objects in locations A1 and A3). We observed elevated levels of *Ccr5* (**Fig. 6B**) and *Ccl5* (**Fig. 6C**) mRNA in the DH of old mice relative to young mice, regardless of behavioral group, though we also detected further increases following behavior in old mice.

**Figure 6.**
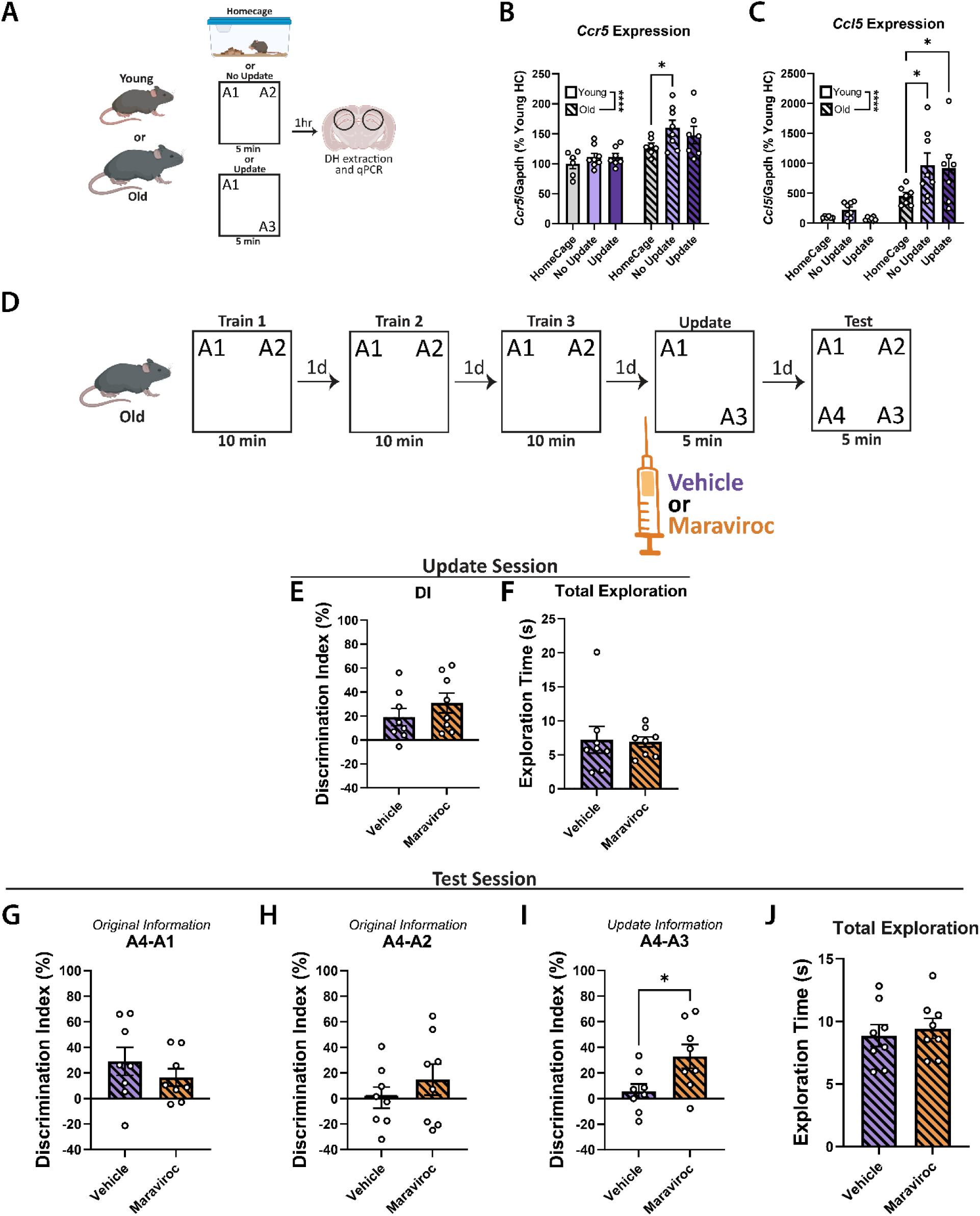
Administration of the CCR5 antagonist maraviroc alleviates age-related memory updating impairments. **A.** Experimental schematic for qPCR. Mice underwent training and were then either left in their homecage, repeated the training session, or underwent the update session. **B.** Old mice had elevated hippocampal expression of *Ccr5* (two-way ANOVA: effect of age: F_(1,37)_ = 23.25, p < 0.0001; *: p < 0.05, Dunnett’s post-hoc test). **C.** Old mice showed elevated hippocampal expression of *Ccl5* (two-way ANOVA: effect of age: F_(1,37)_ = 36.22, p < 0.0001; *: p < 0.05, Dunnett’s post-hoc test). **D.** Experimental schematic for behavior. Old mice were administered either maraviroc or vehicle 1 hr prior to the update session. **E.** At updating, both groups demonstrated intact memory for initial training (one-sample t-tests of A3-A1 DIs against 0: vehicle: t_7_ = 2.63, p = 0.034; maraviroc: t_7_ = 3.72, p = 0.0074), with no differences between groups (unpaired Student’s t-test: t_14_ = 1.07, p = 0.31). **F.** No differences in total exploration time during the update (Mann-Whitney test: U = 26, p = 0.57). **G.** At test, both groups demonstrated intact memory for training position A1 (one-sample t-tests of A4-A1 DIs against 0: vehicle: t_7_ = 2.65, p = 0.033; maraviroc: t_7_ = 2.42, p = 0.046), with no significant differences (unpaired Student’s t-test: t_14_ = 0.98, p = 0.35). **H.** Neither group demonstrated memory for training position A2 (one-sample t-tests of A4-A2 DIs against 0: vehicle: t_7_ = 0.072, p = 0.94; maraviroc: t_7_ = 1.20, p = 0.27), with no group differences (unpaired Student’s t-test: t_14_ = 0.96, p = 0.36). **I.** Only mice in the maraviroc group remembered update position A3 (one-sample t-tests of A4-A1 DIs against 0: vehicle: t_7_ = 0.95, p = 0.37; maraviroc: t_7_ = 3.54, p = 0.0095), performing significantly better than vehicle mice (unpaired Student’s t-test: t_14_ = 2.50, p = 0.025). **J.** There were no differences in exploration time (unpaired Student’s t-test: t_14_ = 0.48, p = 0.64). All data are presented as mean ± SEM. n = 6, 8, 7, 7, 8, 7 (qPCR); 8, 8 (behavior). All male.

Given these age-related changes in *Ccr5* and *Ccl5* mRNA expression, we investigated if pharmacological inhibition of CCR5 would improve memory updating in old mice. Maraviroc is an FDA-approved CCR5 antagonist prescribed for the treatment of HIV,^41^ and prior work has demonstrated that it can improve memory linking in 16 to 18 m.o. mice by mediating engram co-allocation.^26^ However, it has yet to be tested if CCR5 inhibition improves memory updating.

Here, old mice underwent OUL and were administered either maraviroc or vehicle 1 hr prior to the update session (**Fig. 6D**). All mice successfully habituated to the arenas (**Extended Data Fig. 10A**) and explored normally during training (**Extended Data Fig. 10B,C**). During the OUL update, both groups of mice showed robust memory for the training sessions (**Fig. 6E**) and similar patterns of total investigation **(Fig. 6F**).

At the retention test, we observed that both vehicle- and maraviroc-treated animals demonstrated intact memory for location A1 (**Fig. 6G**), though neither group showed memory for location A2 at test (**Fig. 6H**). However, only mice in the maraviroc group demonstrated robust memory for the updated location A3 (**Fig. 6I**), and this updating enhancement was not driven by differences in total investigation time (**Fig. 6J**). Raw percent exploration time for each object during the test session is shown in **Extended Data Fig. 10D**. These results indicate that inhibiting CCR5 ameliorates age-related deficits in memory updating, likely by promoting the co-allocation of the pre- and post-update representations.

## Discussion

The dynamics of neuronal engrams in memory formation and subsequent retrieval are well characterized, but how these ensembles behave during reconsolidation-dependent memory updating remains poorly understood. Here, we demonstrated that dCA1 update engrams in the young brain are preferentially co-allocated to the neuronal ensemble storing the original memory, and that this co-allocation is diminished in the old brain (**Fig. 1**). We demonstrate the relevance of this co-allocation deficit to age-related updating impairments by showing that targeting this deficit through viral manipulations (**Fig. 2**; **Fig. 3**), behavioral interventions (**Fig. 5**) or pharmacology (**Fig. 6**) alleviates age-related deficits in memory updating. Together, these results indicate that engram co-allocation is a common mechanism that underlies many different forms of memory organization, including both memory linking^24,26,27,42^ and updating, and that leveraging recent findings is a promising therapeutic avenue for promoting successful memory updating.

Future work should aim to identify other common mechanisms of memory organization that occur during both memory linking and updating. For instance, we only examined dCA1 here, but we would anticipate that our findings might generalize to other regions where engram co-allocation has been shown to occur during linking, one such additional region being the retrosplenial cortex (RSC).^27^ A recent report indicates that the vmPFC modulates engram co-allocation in the DH during memory linking,^43^ consistent with the role of both of these regions in the generation of memory schema.^44^ Notably, human studies have also implicated the vmPFC during memory updating,^45–47^ further highlighting the importance of top-down control in mediating how memories are locally organized and related to one another in the DH and demonstrating the mechanistic similarities between these two forms of memory organization. Indeed, both memory linking^48^ and memory updating^49^ are also dependent on dopamine release from the locus coeruleus to the DH. Exploring further mechanistic similarities between various forms of memory organization will be useful for identifying mechanisms unique to memory linking or updating and provide a more comprehensive understanding of how the brain organizes and integrates multiple different memories to guide behavior.

In the OUL task, we often observe retroactive interference wherein memory for position A2 (present during the initial training but absent during the update) is impaired in old mice at test,^11^ though this effect does not always occur,^32^ seemingly driven by variability between cohorts. Here, we observed this A2 impairment in old mice during the maraviroc (**Fig. 6**) and STM experiments (**Extended Data Fig. 3**), but not in the TetTag hM3Dq (**Fig. 3**) or the 75-min experiments (**Fig. 5**). Interestingly, it seems that the allocate-and-manipulate DREADD experiment may have specifically alleviated this effect of retroactive interference (**Fig. 2**). This is likely due to the selective activation of hM3Dq-encoding neurons during each training session, thereby strengthening the A2-containing training memory and allowing it to outcompete the interference from the update learning.

Although we demonstrate an important role for dCA1 engram co-allocation in memory updating, we nonetheless did not observe any updating deficits in young mice following selective inhibition of the dCA1 training engram during updating **(Fig. 4; Extended Data Fig. 7**). One possibility is that the young hippocampus may be able to compensate for a decrease in co-allocation by forming a second, update-specific memory that is simply expressed alongside the training memory at test. Thus, co-allocation disruptions might interrupt updating only in old mice due to known age-related increases in susceptibility to memory interference.^50,51^ Alternatively, it may simply be the case that preventing co-allocation is more difficult than inducing it. For instance, the inhibitory effect of the hM4Di DREADD might be insufficient to outcompete the neuronal activation driven by natural retrieval cues. Future experiments could use an inhibitory opsin to investigate if a more carefully timed inhibition of the training engram during updating is sufficient to interfere with this process in young adult mice.

Understanding the mechanisms of memory reconsolidation has been of interest for many years, and many of the molecular and intracellular requirements of this process have been mapped out in detail.^5,6^ Whether these molecular mechanisms occur specifically within engram neurons, however, has yet to be explored. We posit that during memory updating, these established mechanisms of reconsolidation (e.g., protein degradation,^52^ synaptic remodeling,^53^ and *de novo* translation^4^) occur predominately within the re-engaged engram neurons in order to adjust synaptic weights to incorporate new information, though this hypothesis has not yet been tested. Future studies might also explore the molecular processes that drive neurons either into or out of a memory engram during successful memory updating or other forms of ensemble remodeling (see^28^).

One molecular mechanism implicated here in memory updating is the activation of the chemokine receptor CCR5 by its ligand CCL5, as we found that both of these genes are upregulated in the old DH and CCR5 inhibition is sufficient to improve memory updating in old mice (**Fig. 6**). Given the known role of CCR5 in inhibiting neuronal excitability^54^ and in regulating hippocampal engram co-allocation,^26^ it seems that this manipulation improves memory updating in old mice by restoring engram co-allocation during the update session. Interestingly, the only group to exhibit a decrease *Ccl5* mRNA expression relative to homecage was the young adult updating group (**Fig. 6C**), which is also the only group to undergo successful memory updating, suggesting that *Ccl5* may be actively downregulated in the training engram following the update session, thereby permitting the co-allocation necessary for successful memory updating.

Notably, as the CCR5 inhibitor used in these experiments (maraviroc) is already FDA approved for clinical use in humans, our findings indicate this drug may have additional applications in managing age-related cognitive decline. Future research should investigate if maraviroc facilitates other forms of memory updating (e.g., of fearful memories or in the context of addiction) and if these observations translate to human patients. Further exploration here can inform future therapeutics intended to treat age-related cognitive decline as well as psychiatric disorders characterized by persistent and maladaptive memories, for instance PTSD and OCD.

## Methods

### Subjects

All mice used for these experiments were C57BL/6J. Young adult mice were between 3 and 4 months of age at the start of experiments and were acquired from the Jackson Laboratory while old mice were between 18 and 20 months of age at the start of experiments and were acquired from the NIA Aged Rodent Colony (maintained by Charles River, Raleigh, NC). Mice were housed either individually (for engram tagging experiments) or in groups of four (other experiments) in a temperature- and humidity-controlled environment with *ad libitum* access to food and water and a 12-hour light/dark cycle (lights on at 6:00 AM). All experiments were performed during the light phase of the cycle. Mice were randomly assigned to groups and appropriately counterbalanced. All experiments were approved by the Institutional Animal Care and Use Committee at the Pennsylvania State University.

### Objects in Updated Locations (OUL)

OUL was performed as previously described.^11,29,32^ In brief, mice were handled for four days in the behavior room (2 min each day) and then habituated to the polypropylene arenas filled with bedding (23.0 cm x 30.0 cm x 23.0 cm) for 5 min each day for six consecutive days. Following habituation, mice underwent three consecutive days of 10-min training in which identical objects (glass 200mL tall-form beakers filled with hydraulic cement) were presented in locations A1 and A2 of the arena. This 3-day training protocol allows old mice to successfully learn the training prior to the update^11^ and was used for both age groups to maintain consistency. Twenty-four hours following the last training, mice underwent the 5-min update session with objects presented in locations A1 and A3. Finally, mice were given a 5-min retention test the next day in which objects were presented in all previously seen locations (A1, A2, and A3) and in the novel location A4. The orientations of these locations were counterbalanced as previously described.^29^ Deviations from this general protocol are described on an experiment-by-experiment basis in the Results.

Movement data were calculated using Ethovision (Noldus, Leesburg, VA) to verify that mice habituated to the OUL context by exhibiting a decrease in distance moved (**Extended Data Fig. 2D, 3H, 4A, 5A, 6A, 7I, 9A, 9E, 10A**) prior to beginning the training sessions. Object investigation was determined by manually scoring behavioral videos on a frame-by-frame basis by a blinded investigator using the GUI of DeepEthoGram.^55^ Investigation was defined as any frame in which the mouse had all four paws on the ground with its nose pointed directly at the object and within 1 cm of the object and was not otherwise biting, climbing, digging, or rearing on the object.^56^ Discrimination indices (DIs) were calculated to quantify memory using the general formula DI = (t_n_ – t_f_)/(t_n_ + t_f_) × 100% where t_n_ is the time spent investigating an object in a novel location and t_f_ is the time spent investigating an object in a familiar location. For the OUL test session, three separate DIs were calculated per animal, each comparing the novel location A4 against a different familiar location (A1, A2, or A3). Preference for A4 over A1 or A2 was interpreted as memory for the initial training, while preference for A4 over A3 was interpreted as memory for the update. DIs were additionally calculated during each training session, to verify animals exhibited no innate preference for a particular side of the arena, and during the update session, to confirm animals successfully learned the initial training. Any animal that failed to explore all four objects during the test session, investigated for less than three seconds total during the test session, exhibited a DI greater than 20 or less than -20 on all three training days, exhibited a DI less than -20 during the update session (other than during the 1-min subthreshold update), or exhibited a DI more than 2 standard deviations away from their group mean was excluded from all analyses.

### Adeno-associated virus (AAV) Administration

AAV9-cFos-tTA (CV17158-AV9; 1.1 × 10^14^ GC/mL) was purchased from Charles River. AAV9-TRE-EGFP (VB220405-1200ren; 4.5 × 10^13^ GC/mL), AAV9-TRE-hM3Dq-EGFP (VB230804-1278pfr; 1.4 × 10^13^ GC/mL), and AAV9-TRE-hM4Di-EGFP (VB241118-1271dvc; 7.6 × 10^13^ GC/mL) were cloned and packaged by Vectorbuilder. AAV9-hSyn-hM3Dq-mCherry (Addgene plasmid #50474; 2.9 × 10^13^ GC/mL, diluted to 5.0 × 10^10^ GC/mL) and AAV9-hSyn-mCherry (Addgene plasmid #114472; 2.4 × 10^13^ GC/mL, diluted to 5.0 × 10^10^ GC/mL) were gifts via Addgene from Bryan Roth and Karl Deisseroth respectively.

AAVs were administered bilaterally to the DH via stereotaxic surgery as previously described.^57,58^ Briefly, mice were anesthetized with isoflurane in oxygen (induced at 3%, maintained at 1-2%, v/v) and placed in the stereotaxic head frame, and then a craniotomy was performed with a micromotor drill. Injection needles were lowered at a rate of 0.2 mm/15 sec to the final coordinates (from Bregma) of AP: -2.00 mm; ML: ±1.50 mm; and DV: -1.50 mm. After a 2-min pause, 0.4 μL of virus (or viral cocktail) was infused at a rate of 6 μL/hr. The injectors were left in place for 5 min after the completion of the viral infusion, after which they were raised by 0.1 mm and allowed to rest for an additional 2 min. Finally, injectors were withdrawn from the injection site at a rate of 0.1 mm/15 sec. Animals were administered 5 mg/kg subcutaneous meloxicam as an analgesic, given 0.5 mL sterile saline for hydration, and allowed to recover in a clean, heated cage. All injections were performed at least two weeks prior to behavioral experiments.

### Engram Tagging

Memory-encoding engrams were labeled using a viral TetTag approach.^23,30^ Specifically, AAV9-cFos-tTA; one of AAV9-TRE-EGFP, AAV9-TRE-hM3Dq-EGFP, or AAV9-TRE-hM4Di-EGFP; and sterile PBS were combined 1:1:2, and 0.4 μL of the resultant cocktail was administered bilaterally to dCA1. The final titers used (after combining solutions) were 2.75 × 10^13^ for AAV9-cFos-tTA, 1.13 × 10^13^ for AAV9-TRE-EGFP, 3.5 × 10^12^ for AAV9-TRE-hM3Dq-EGFP, and 1.9 × 10^13^ for AAV9-TRE-hM4Di-EGFP. Animals were placed on a 200 mg/kg Doxycycline (Dox) diet (S3888; Bio-Serv, Flemington, NJ) at least one week prior to stereotaxic surgery and maintained on this diet throughout the experiment. Forty-eight hours prior to tagging, animals were taken off Dox, given control feed (5053; LabDiet, St. Louis, MO), and left undisturbed until the first day of OUL training. Immediately following this training session, animals were placed back on Dox. We have demonstrated that these TetTag constructs are both activity- and Dox-dependent in our hands, and that an off Dox period of 48 hrs induces robust GFP labelling in dCA1 (**Extended Data Fig. 1**).

### Clozapine-n-oxide Injections

To activate DREADDs, clozapine-n-oxide (CNO) was administered via intraperitoneal injection. In DREADD experiments, mice were manually restrained daily for four consecutive days (following habituation) to acclimate them to the injection procedure. Stock solutions of CNO (4936; Tocris Bioscience) were prepared in DMSO at a concentration of 30 mg/mL. On the morning of the injection day, these stocks were diluted in sterile saline to a final concentration of 0.3 mg/mL and then administered intraperitoneally 30 min prior to the behavioral session at a final dose of 3 mg/kg, based on previous work.^58,59^ For the hM4Di replication experiment in young mice verifying that inhibition of the training engram does not disrupt memory updating (**Extended Data Fig. 7**), the CNO dose was increased to 5 mg/kg to ensure that our null effect was not due to an insufficient CNO dose.

### Maraviroc Injections

For the maraviroc experiment, mice were manually restrained daily for four consecutive days (following habituation) to acclimate them to the injection procedure. On the update day, maraviroc (Selleck Chemicals, Houston, TX) was dissolved in sterile saline containing 10% DMSO to a concentration of 2 mg/mL and administered intraperitoneally 1 hr prior to the update session at a final dose of 20 mg/kg, chosen based on previous work.^26,60^

### Immunohistochemistry and Cell Counting

Animals were euthanized via cervical dislocation 60 min after behavior and decapitated with surgical scissors. Brains were removed from the skull and post-fixed in 4% paraformaldehyde (Fisher Scientific, Waltham, MA) at 4 °C for 16-18 hrs. After post-fixing, brains were washed in PBS, then cryoprotected in PBS with 15% sucrose for 12 hrs followed by PBS with 30% sucrose and 0.01% sodium azide for at least three days. After cryoprotection, brains were sectioned with a Leica CM1950 Cryostat (Leica Biosystems, Wetzlar, Germany). Coronal sections of 50 μm were collected from the dorsal hippocampus and stored at 4 °C in PBS with 0.01% sodium azide until staining. Sections were blocked in blocking buffer (PBS with 0.5% Triton-X and 5% NGS) for 1 hr at RT. Then, sections were incubated in blocking buffer with primary antibody for 24 hrs at 4 °C. The next day, sections were washed with PBS with 0.5% Triton-X and then incubated with secondary antibody in blocking buffer for 2 hrs at RT. Sections were again washed with PBS with 0.5% Triton-X and then incubated with DAPI (1:10000) for 15 min, given a final wash with PBS, and then mounted on SuperFrost Plus Slides (Fisher Scientific, Waltham, MA) in Vectashield Plus (Vector Labs, Plain City, OH) before being sealed with nail polish (Black Onyx; OPI Products, Calabasas, CA). Antibodies used were: Chicken anti-GFP (1:1000, A10262; ThermoFisher, Frederick, MD), Rabbit anti-cFos (1:2000, 226003; Synpatic Systems, Gottingen, Germany), Rabbit anti-mCherry (1:500, ab167453; Abcam, Waltham, MA), Alexa 488 Goat anti-chicken (1:500, A11039; ThermoFisher, Frederick, MD), and Alexa 555 Goat anti-Rabbit (1:500, A21428; ThermoFisher, Frederick, MD).

For each mouse, a single z-stack was taken per hemisphere of up to three separate dCA1 sections (4-6 stacks per mouse) using a STELLARIS 5 white light laser confocal microscope (Leica Biosystems, Wetzlar, Germany) with a 20X objective. Microscope settings (zoom, exposure time, laser strength, gain, etc.) were kept identical within each experiment. Using the Fiji package of ImageJ (NIH, Bethesda, MD), a blinded investigator manually counted the number of GFP^+^, c-Fos^+^, and GFP^+^c-Fos^+^ cells in dCA1 of each stack. Measurements from each stack were then pooled per mouse to generate average values that were used for subsequent analysis. Within each experiment, all exposure and brightness/contrast settings were kept consistent. The top and bottom of each stack was ignored to minimize the effects of edge artifacts and prevent the counting of partial cells. For each stack, the total number of DAPI cells within dCA1 was estimated using the Voronoi Threshold Labeler of the BioVoxxel 3D Box package for Fiji.^61^ Reactivation Rates were calculated as (GFP^+^c-Fos^+^)/(GFP^+^) × 100%. Similarity Indices were calculated as (GFP^+^ c-Fos^+^)/((GFP^+^) + (c-Fos^+^)-(GFP^+^ c-Fos^+^)) × 100%, as previously described.^33,34^

### qPCR

Animals were euthanized via cervical dislocation 60min after behavior and decapitated with surgical scissors. Brains were removed from the skull and flash-frozen in 2-methylbutane (Fisher Scientific, Waltham, MA). Brains were stored at -80 °C before being sectioned with a Leica CM1950 Cryostat (Leica Biosystems, Wetzlar, Germany). Punches of 500 μm were collected from the dorsal hippocampus and stored at -80 °C prior to RNA extraction. RNA was extracted from punches with RNeasy Mini Kits (Qiagen, Germantown, MD) and cDNA was generated with High-capacity cDNA Reverse Transcription Kits (ThermoFisher, Frederick, MD). PrimeTime primer/probe assays were generated using the IDT PrimerQuest Design Tool (IDT, Coralville, IA) and used to quantify the expression of *Ccr5*, *Ccl5*, and *Gapdh*. Exact sequences are presented in **Table S1**. Proprietary Roche algorithms were used to quantify gene expression, with *Gapdh* serving as the reference assay.

### Electrophysiology

Mice were deeply anesthetized via inhaled isoflurane (5% in oxygen, v/v) and rapidly decapitated. Brains were quickly removed and processed according to the N-methyl-D-glucamine (NMDG) protective recovery method.^62^ Brains were immediately placed in ice-cold oxygenated NMDG-HEPES artificial cerebrospinal fluid (aCSF) containing the following, in mM: 92 NMDG, 2.5 KCl, 1.25 NaH_2_PO_4_, 30 NaHCO_3_, 20 HEPES, 25 glucose, 2 thiourea, 5 Na-ascorbate, 3 Na-pyruvate, 0.5 CaCl_2_ 2H_2_O, and 10 MgSO_4_ 7H_2_O (pH’d to 7.3–7.4). The hippocampus was identified according to the Allen Mouse Brain Atlas. 300-μm coronal slices containing the hippocampus were prepared on a Compresstome Vibrating Microtome VF-300–0Z (Precisionary Instruments, Greenville, NC), and transferred to heated (31°C) NMDG-HEPES (in mM: 124 NaCl, 4.4 KCl, 2 CaCl_2_, 1.2 MgSO_4_, 1 NaH_2_PO_4_, 10.0 glucose, and 26.0 NaHCO_3_, pH 7.4, mOsm 300– 310), for a maximum of 10 min. Slices were then transferred to heated (31°C) oxygenated normal aCSF where they were allowed to rest for at least 1 hr before use. Finally, slices were moved to a submerged recording chamber where they were continuously perfused with the recording aCSF (2 mL per min flow rate, 31°C). Recording electrodes (3–6 MΩ) were pulled from thin-walled borosilicate glass capillaries with a Narishige PC-100 Puller. A potassium-gluconate (KGluc)-based intracellular recording solution, containing the following (in mM): 135 K-Gluc, 5 NaCl, 2 MgCl_2_, 10 HEPES, 0.6 EGTA, 4 Na_2_ATP, and 0.4 Na_2_GTP (287–290 mOsm, pH 7.35) was used. Cells were identified by the presence of GFP and were patched in the dCA1 region of the hippocampus. Following rupture of the cell membrane, cells were held in current-clamp. A minimum of 5 min stable baseline was acquired prior to experiments and bath application. Following a 5 min stable baseline 10µM clozapine N-oxide (CNO) water soluble (Hello Bio HB187) was bath applied. Resting membrane potential (RMP) was recorded during the entire 10 min CNO application period. Experiments were conducted blinded to DREADD condition, and mice were randomized between groups and experiment day. Slices were replaced after CNO application to ensure neurons did not receive CNO application more than once. Input resistance was monitored intermittently throughout each experiment, and when it deviated by more than 20% the experiment was discarded. A single mouse was excluded from the hM4Di group, as both cells from this animal depolarized in response to CNO administration.

### Statistics

Data are represented as Mean ± SEM in all figures. Sample sizes were chosen on the basis of previous studies^11,63^ but no statistical method was used to predetermine sample size. All analyses were conducted in GraphPad Prism 10 (GraphPad Software, San Diego, CA). Data normality was verified with the Shapiro-Wilk test and variances between groups were checked with an F test prior to subsequent analysis. Normally distributed data were analyzed using unpaired Student’s t-tests (Welch’s t-tests when variances were significantly different) or ANOVAs followed by Dunnett’s post-hoc tests. Data that were not normally distributed were analyzed using Mann-Whitney U tests. For all statistical tests, α was set to 0.05 and all comparisons were two-tailed. For experiments using male and female mice, data were preliminarily tested for an effect of sex. However, as no sex differences were detected in these experiments, male and female mice were pooled for subsequent analyses to increase power.

### Data Availability

The data supporting the findings of this study are uploaded to the Inter-university Consortium for Political and Social Research (ICPSR). Additional data files and materials are available from the corresponding author upon reasonable request.

## Acknowledgements

We would like to thank all members of the Kwapis Lab for scientific discussion and technical assistance. We also thank Aaron Fleischer and Steve Ramirez for sharing TetTag constructs and protocols and Jan Brocher for help with automated three-dimensional cell quantification. Additionally, we thank Melissa Sharpe and Moriel Zelikowsky for helpful comments on an earlier version of this manuscript. Figures were made with BioRender. This work was supported by National Institutes of Health grants F31AG087533 (C.A.B.), R01AG07041 (J.L.K.), RO1AA031472 (N.A.C.), and R01AA029403 (N.A.C.); by the McKnight Brain Research Foundation/AFAR Innovator Award in Cognitive Aging and Memory Loss (J.L.K.); and by the Hevolution/AFAR New Investigator Award in Aging Biology and Geroscience Research (J.L.K.).

## Author Contributions

C.A.B. and J.L.K. designed behavioral, molecular, and immunohistochemical experiments. C.A.B., A.G.D, T.A.W., D.J.B., A.R.M., and C.R.M. conducted behavioral, molecular, and immunohistochemical experiments. A.R.S., D.F.B., and N.A.C. designed, conducted, and analyzed electrophysiological experiments. S.M. contributed to the design and validation of plasmids and viral constructs. G.C.P. and C.W.S. contributed to data collection. C.A.B., A.G.D., and J.L.K. analyzed and interpreted the results. C.A.B. and J.L.K. wrote and revised the manuscript. All authors approved the manuscript.

## Supplementary Materials

**Extended Data Figure 1.**
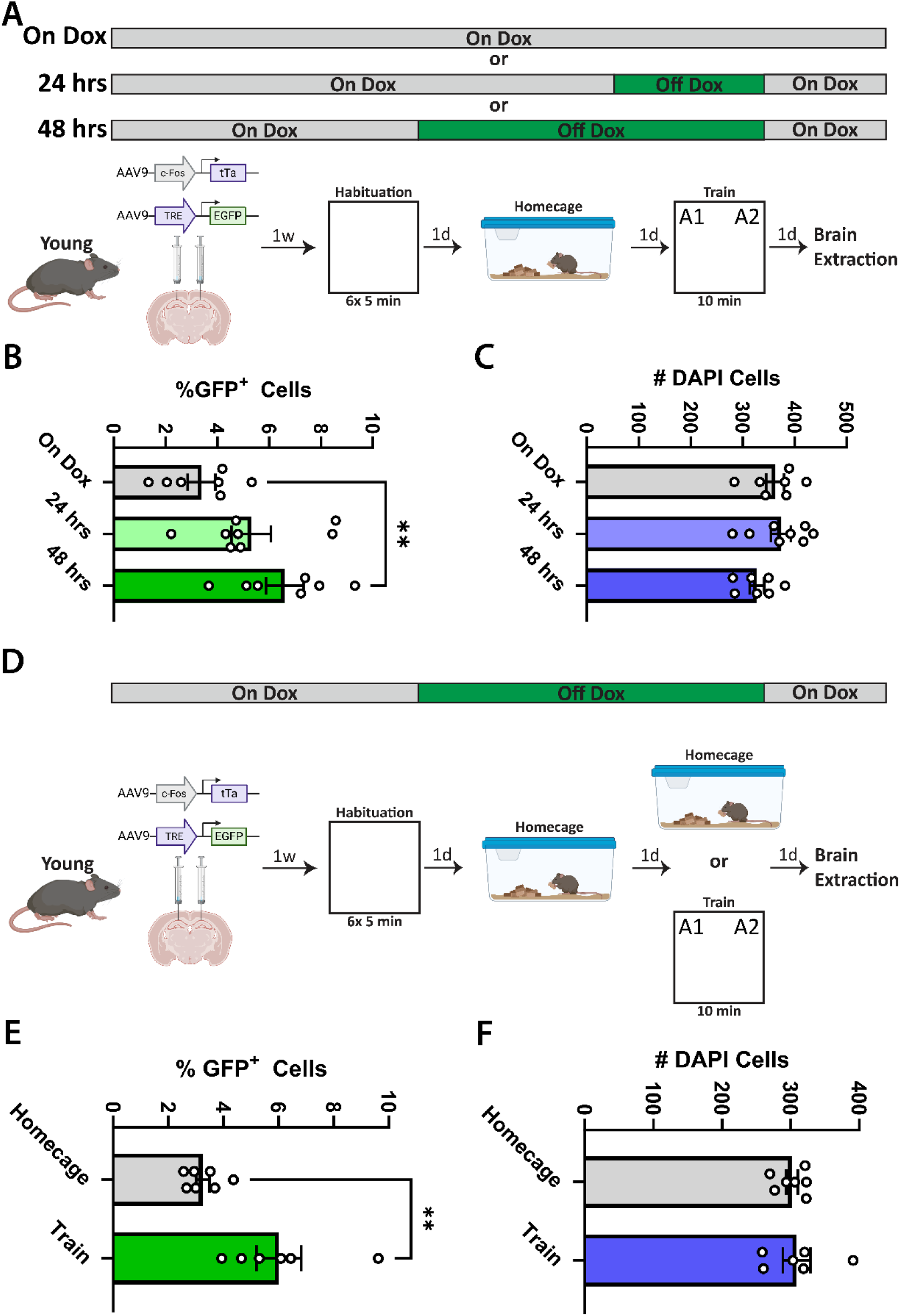
Validations of the dual AAV TetTag labelling system. **A.** Experimental schematic for the Dox-dependency validation. Young mice were administered dCA1 injections of the viral TetTag system and then, 2 wks later, were taken off Dox either 24 or 48 hrs prior to OUL training. Following training, Dox was restored and mice were sacrificed 24 hrs later and the percentage of GFP^+^ cells in dCA1 was assessed. **B.** Significantly more GFP^+^ cells were detected in dCA1 in mice that were off Dox for 48 hrs compared to those left on Dox (one-way ANOVA of %GFP^+^ cells, F_(2, 19)_ = 5.26, p = 0.015. ** denotes p < 0.01 on Dunnett’s multiple comparisons test against On Dox group). **C.** No difference was detected between groups in the number of DAPI cells per stack (one-way ANOVA of #DAPI cells, F_(2, 19)_ = 1.95, p = 0.17). **D.** Experimental schematic for the activity-dependency validation. Young mice were administered dCA1 injections of the viral TetTag system and then, 2 wks later, were taken off Dox and underwent either an OUL training session or were left in their homecage. Following training, Dox was restored and mice were sacrificed 24 hrs later and the percentage of GFP^+^ cells in dCA1 was assessed. **E.** Significantly more GFP^+^ cells were detected in the train group relative to the homecage group (Welch’s t-test: t_5.90_ = 3.26, p = 0.018). **F.** No difference was detected between groups in the number of DAPI cells per stack (unpaired Student’s t-test: t_11_ = 0.34, p = 0.74). All data are presented as mean ± SEM.

**Extended Data Figure 2.**
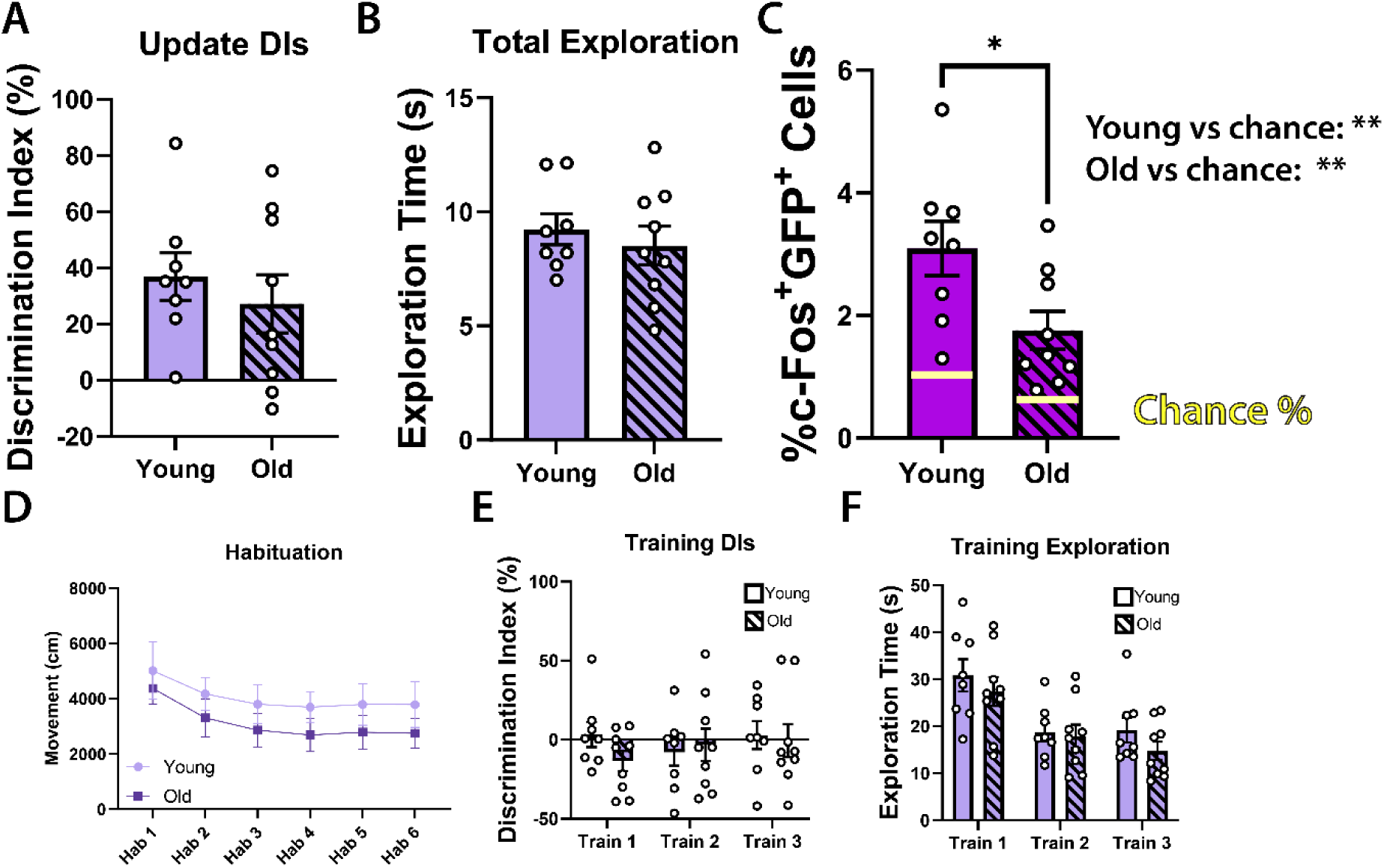
Additional data from the engram co-allocation experiment (Figure 1). **A.** DIs from the update session. Both groups of mice demonstrated intact memory for the initial training during the update (one-sample t-tests of A3-A1 DIs against 0: young: t_7_ = 4.37, p = 0.0033; old: t_8_ = 2.63, p = 0.030), with no differences detected between groups (unpaired Student’s t-test: t_15_ = 0.72, p = 0.49). **B.** No differences were detected in total investigation time during the update session (unpaired Student’s t-test: t_15_ = 0.64, p = 0.53). **C.** Young mice had a significantly higher proportion of c-Fos^+^GFP^+^ cells in dCA1 than old mice (unpaired Student’s t-test: t_15_ = 2.51, p = 0.024), though both groups exhibited more than would be expected by chance (one-sample t-tests of %c-Fos^+^GFP^+^ against chance: young: t_7_ = 4.50, p = 0.0028; old: t_8_ =3.37, p = 0.0098). **D.** Both groups habituated normally to the arenas, though old mice moved significantly less, in-line with physical symptoms of aging (two-way RM ANOVA: main effects of session, age, and subject). **E.** Neither group had a significant object preference during the initial training sessions (two-way RM ANOVA, no significant effects). **F.** Both groups spent comparable time investigating objects during training (two-way RM ANOVA, main effects of session and subject). All data are presented as mean ± SEM.

**Extended Data Figure 3.**
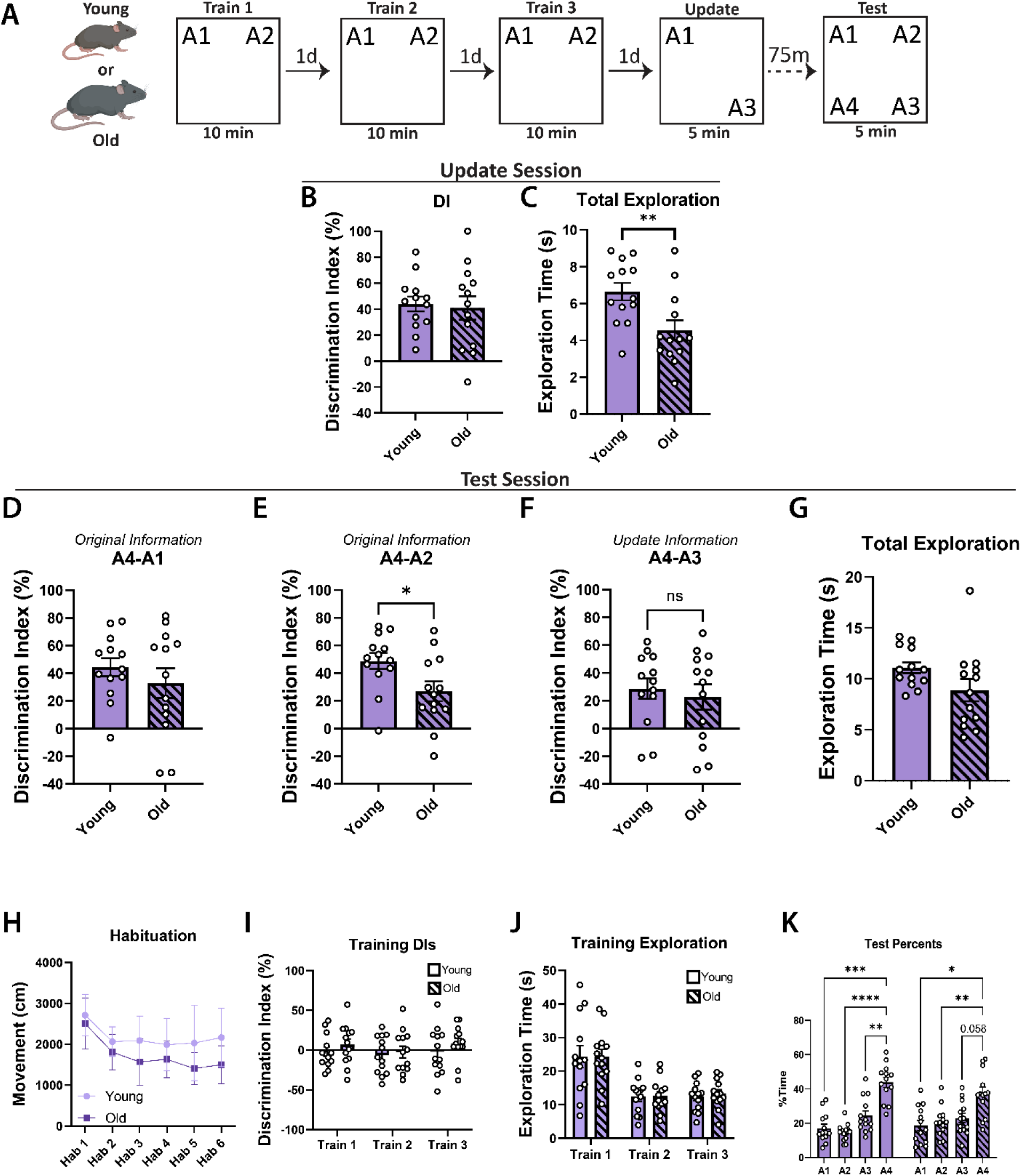
Age-related updating deficits are specific to long-term memory and not observable at an STM test. **A.** Experimental schematic. Young and old mice underwent OUL and were tested 75 min after the update session. **B.** During the update session, both young and old mice demonstrated intact memory for the initial training (one-sample t-tests of A3-A1 DIs against 0: young: t_12_ = 7.65, p < 0.0001; old: t_12_ =4.50, p = 0.0007), with no differences between groups (unpaired Student’s t-test: t_24_ = 0.29, p = 0.78). **C.** Young mice spent significantly more time investigating during the update session (unpaired Student’s t-test: t_24_ = 2.89, p = 0.0080). **D.** At test, both groups remembered training location A1 (one-sample t-tests of A4-A1 DIs against 0: young: t_12_ = 6.80, p < 0.0001; old: t_12_ = 3.07, p = 0.0097), with no differences between groups (unpaired Student’s t-test: t_24_ = 0.92, p = 0.37). **E.** Both groups remembered training location A2 (one-sample t-tests of A4-A2 DIs against 0: young: t_12_ = 8.43, p < 0.0001; old: t_12_ = 3.74, p = 0.0028), though young mice exhibited significantly better memory for this location (unpaired Student’s t-test: t_24_ = 2.39, p = 0.025), reflecting age-related susceptibility to memory interference. **F.** Both young and old mice remembered the update position A3 (one-sample t-tests of A4-A3 DIs against 0: young: t_12_ = 3. 88, p = 0.0022; old: t_12_ = 2.46, p = 0.030), with no differences between groups (unpaired Student’s t-test: t_24_ = 0.51, p = 0.61). **G.** No differences in total investigation time were observed at test, though young mice were trending higher (Mann-Whitney test: U = 49.5, p = 0.074). **H.** Both groups habituated to the arenas as expected, with young mice moving more overall than old mice (two-way RM ANOVA: main effects of session, age, and subject). **I.** Neither group had a significant object preference during the training sessions (two-way RM ANOVA, significant effect only of subject). **J.** Both groups spent comparable time investigating objects during training (two-way RM ANOVA, main effects of session and subject). **K.** Investigation times at test displayed as percentages (two-way RM ANOVA, main effect of object; * denotes p < 0.05, ** denotes p < 0.01, *** denotes p < 0.001, **** denotes p < 0.0001 on Dunnett’s multiple comparisons test against A4). All data are presented as mean ± SEM. n = 13, 13, all male.

**Extended Data Figure 4.**
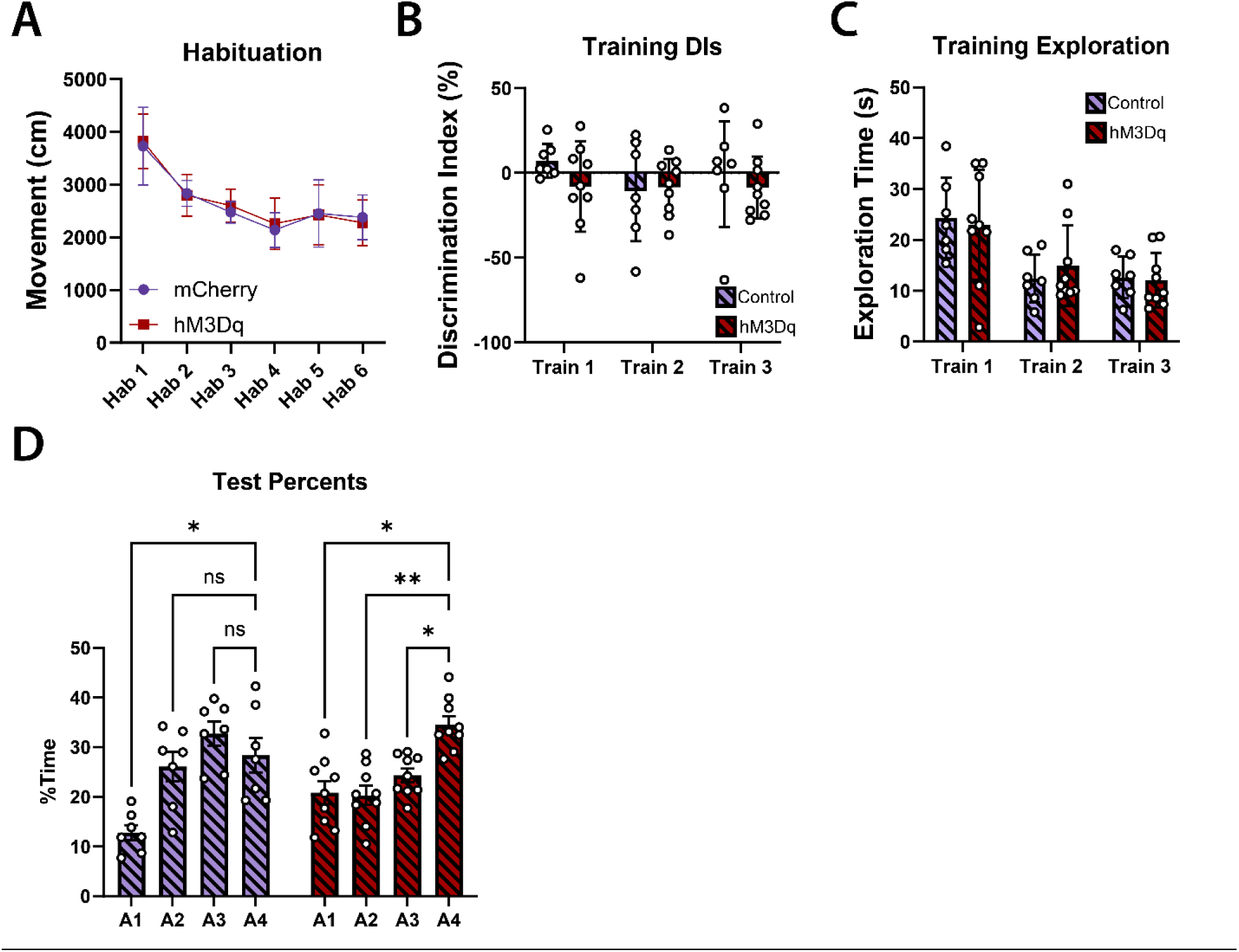
Additional data from the allocate-and-manipulate hM3Dq experiment (Figure 2). **A.** Both groups habituated to the arenas as expected (two-way RM ANOVA: main effects of session and subject). **B.** Neither group had a significant object preference during the training sessions (two-way RM ANOVA, no significant effects). **C.** Both groups spent comparable time investigating objects during training (two-way RM ANOVA, main effect only of session). **D.** Investigation times at test displayed as percentages (two-way RM ANOVA, effect of object and object x virus interaction; * denotes p < 0.05, ** denotes p < 0.01, *** denotes p < 0.001, **** denotes p < 0.0001 on Dunnett’s multiple comparisons test against A4). All data are presented as mean ± SEM.

**Extended Data Figure 5.**
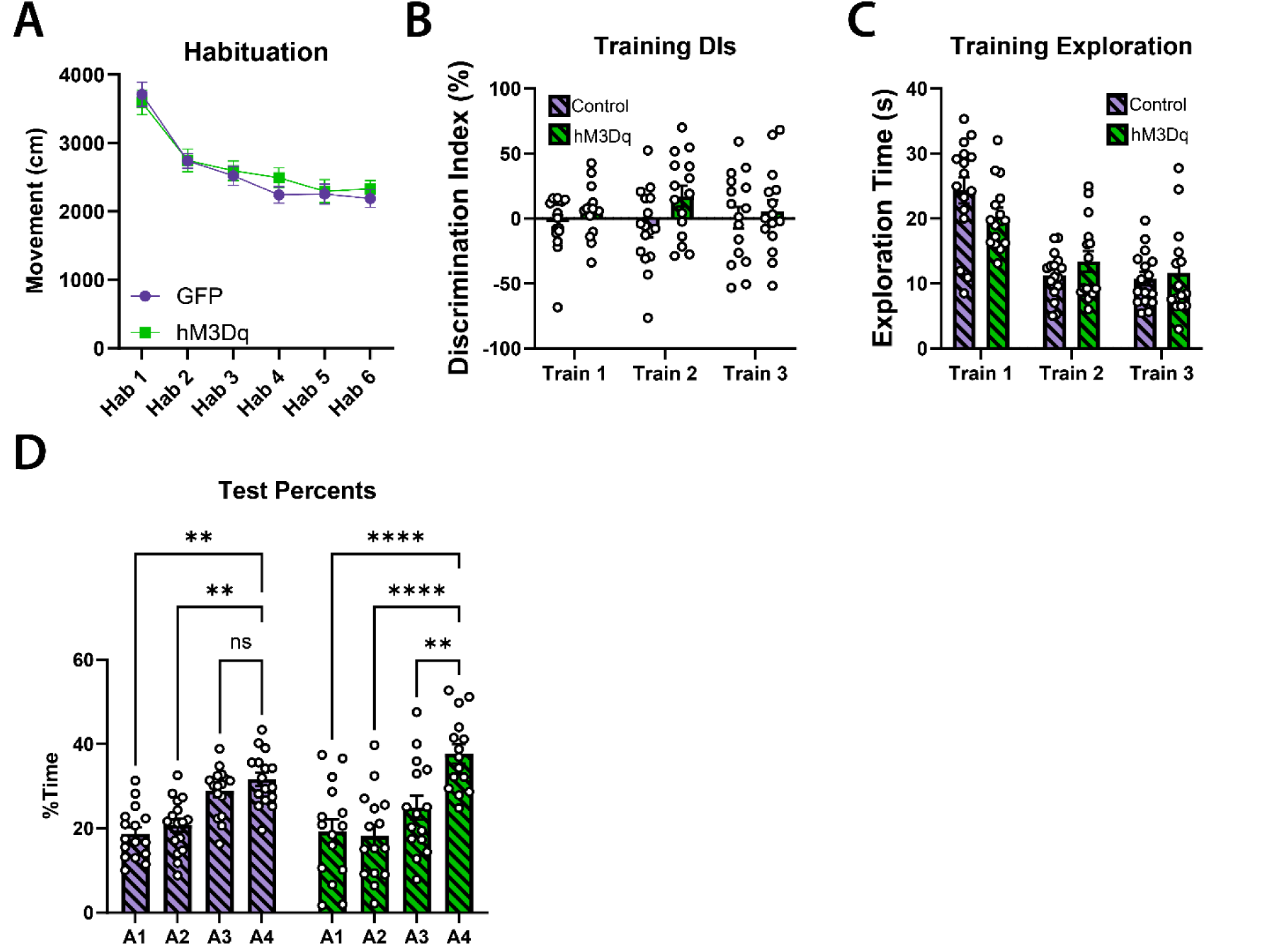
Additional data from the TetTag hM3Dq experiment (Figure 3). **A.** Both groups habituated to the arenas as expected (two-way RM ANOVA: main effects of session and subject). **B.** Neither group had a significant object preference during the training sessions (two-way RM ANOVA, no significant effects). **C.** Both groups spent comparable time investigating objects during training, though there was a significant interaction between the virus and the session (two-way RM ANOVA, effects of session, subject, and session x virus interaction). **D.** Investigation times at test displayed as percentages (two-way RM ANOVA, effect of object; * denotes p < 0.05, ** denotes p < 0.01, *** denotes p < 0.001, **** denotes p < 0.0001 on Dunnett’s multiple comparisons test against A4). All data are presented as mean ± SEM.

**Extended Data Figure 6.**
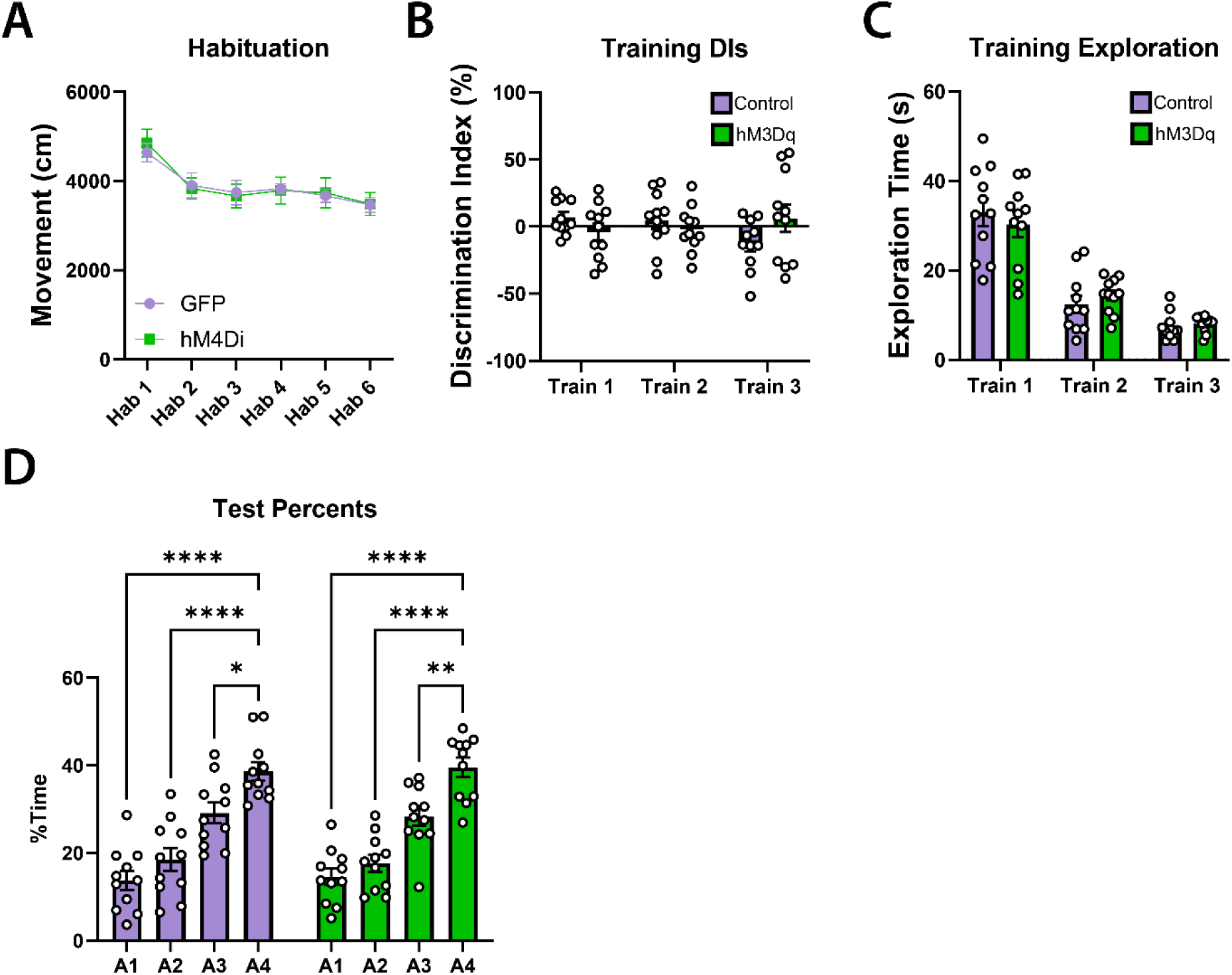
Additional data from the TetTag hM4Di experiment (Figure 4). **A.** Both groups habituated to the arenas as expected (two-way RM ANOVA: main effects of session and subject). **B.** Neither group had a significant object preference during the training sessions (two-way RM ANOVA, no significant effects). **C.** Both groups spent comparable time investigating objects during training (two-way RM ANOVA, effect of session). **D.** Investigation times at test displayed as percentages (two-way RM ANOVA, effect of object; * denotes p < 0.05, ** denotes p < 0.01, *** denotes p < 0.001, **** denotes p < 0.0001 on Dunnett’s multiple comparisons test against A4). All data are presented as mean ± SEM.

**Extended Data Figure 7.**
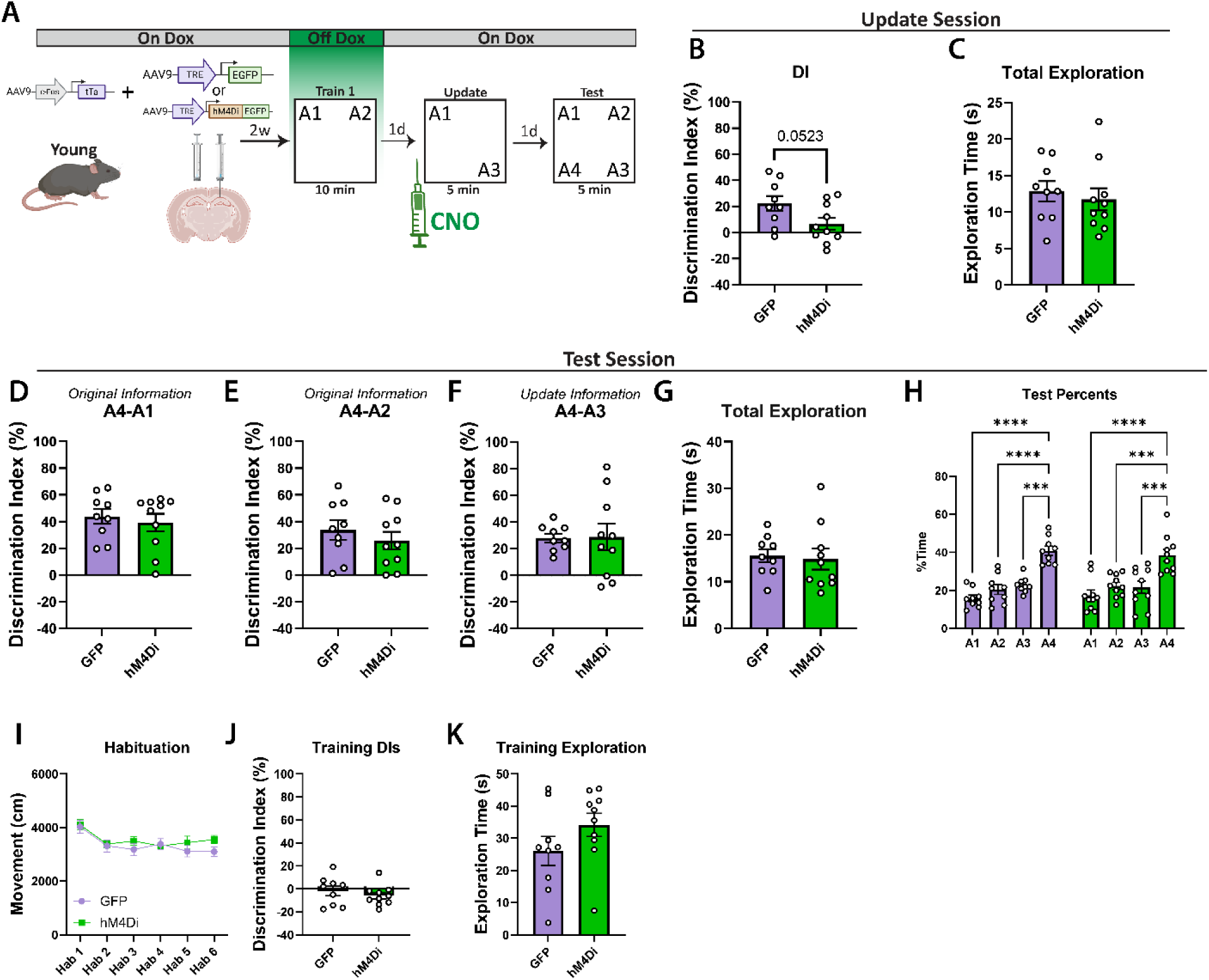
Replication of the TetTag hM4Di experiment (Figure 4) with a single training session and an increased CNO dose. **A.** Experimental schematic. Animals were injected with the dual virus TetTag system (expressing either hM4Di-GFP or GFP alone) two weeks prior to engram tagging. Mice were removed from Dox 48 hrs prior to training to label the training engram with GFP and were then promptly restored to Dox. CNO was administered 30 min prior to the update session. This experiment differed from that in Figure 4 in that only a single day of training was used and a higher CNO dose (5 mg/kg) was administered. **B.** During the update session, only the GFP group demonstrated intact memory for the initial training (one-sample t-tests of A3-A1 DIs against 0: GFP: t_8_ = 3.83, p = 0.0050; hM4Di: t_9_ = 1.39, p = 0.20), and the hM4Di group tended to perform worse than the GFP group (unpaired Student’s t-test: t_17_ = 2.09, p = 0.052), consistent with inactivation of the training engram. **C.** No differences were observed in object investigation during the update session (unpaired Student’s t-test: t_17_ = 0.53, p = 0.60). **D.** At test, both groups of mice demonstrated intact memory for training position A1 (one-sample t-tests of A4-A1 DIs against 0: GFP: t_8_ = 7.79, p < 0.0001; hM4Di: t_9_ = 6.04, p = 0.0002) with no differences observed between groups (Mann-Whitney test: U = 42, p = 0.84). **E.** Both groups also remembered position A2 from training (one-sample t-tests of A4-A2 DIs against 0: GFP: t_8_ = 4.59, p = 0.0018; hM4Di: t_9_ = 3.83, p = 0.0041), with no differences detected between groups (unpaired Student’s t-test: t_17_ = 0.82, p = 0.43). **F.** Both groups of mice also demonstrated intact memory for update position A3 (one-sample t-tests of A4-A3 DIs against 0: GFP: t_8_ = 8.65, p < 0.001; hM4Di: t_9_ = 2.94, p = 0.017), with no differences detected between groups (Welch’s t-test: t_10.93_ = 0.079, p = 0.94). **G.** We additionally detected no differences in total investigation time at test (unpaired Student’s t-test: t_17_ = 0.26, p = 0.80). **H.** Investigation times at test displayed as percentages (two-way RM ANOVA, effect of object; * denotes p < 0.05, ** denotes p < 0.01, *** denotes p < 0.001, **** denotes p < 0.0001 on Dunnett’s multiple comparisons test against A4). **I.** Both groups habituated to the arenas as expected (two-way RM ANOVA: main effects of session and subject). **J.** Neither group had a significant object preference during training (unpaired Student’s t-test: t_17_ = 0.83, p = 0.42). **K.** Both groups spent comparable time investigating objects during training (unpaired Student’s t-test: t_17_ = 1.42, p = 0.17). All data are presented as mean ± SEM. n = 9 (3F), 10 (4F).

**Extended Data Figure 8.**
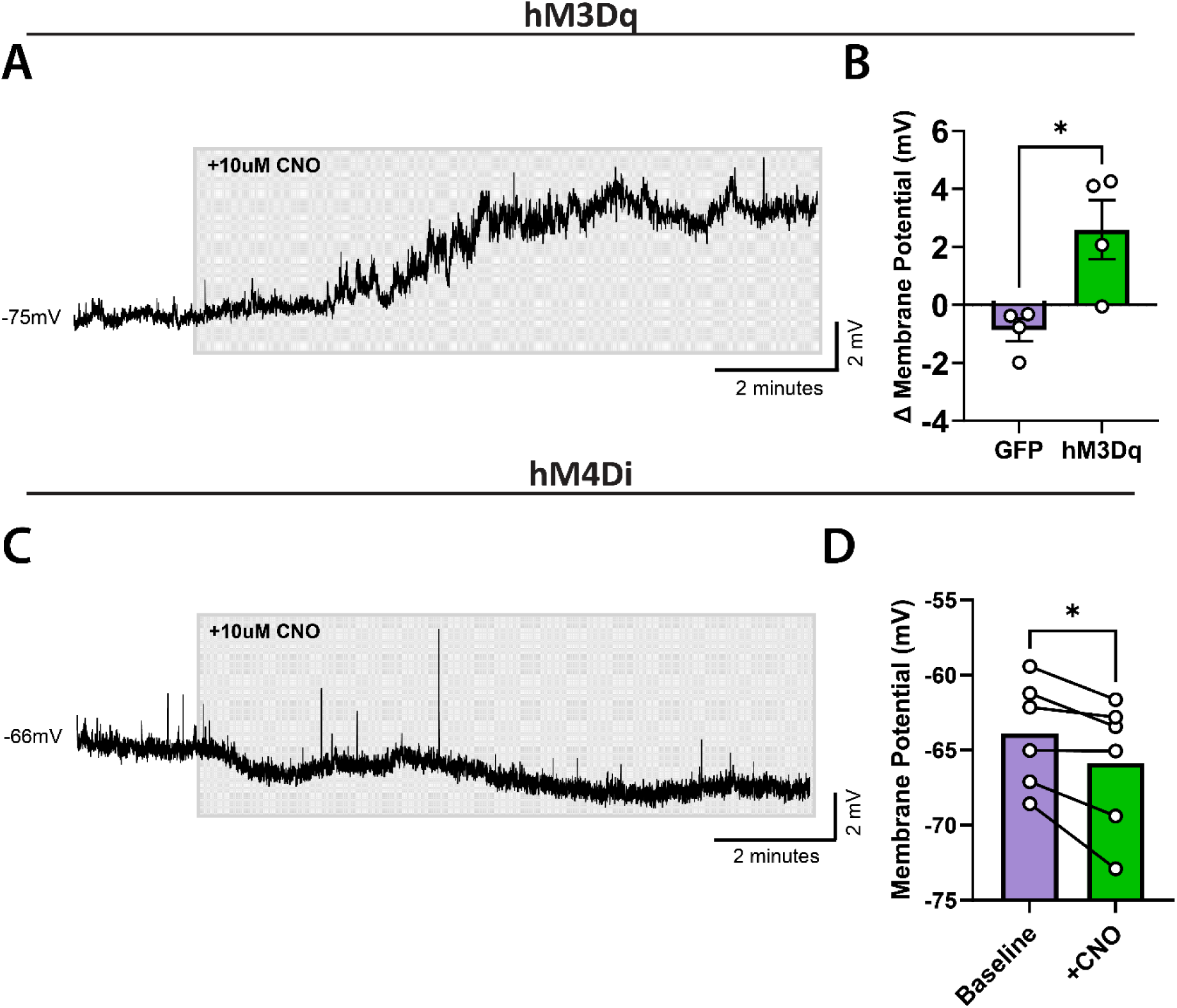
Electrophysiological validation of hM3Dq and hM4Di DREADDs. **A.** Resting membrane potential of a representative hM3Dq^+^ cell before and during CNO administration. **B.** Administration of CNO increased the membrane potential of hM3Dq^+^ cells relative to GFP^+^ controls (unpaired Student’s t-test: t_6_ = 3.19, p = 0.019). **C.** Resting membrane potential of a representative hM4Di^+^ cell before and during CNO administration. **D.** CNO significantly decreased the membrane potential of hM4Di^+^ cells relative to pre-CNO baseline (paired Student’s t-test: t_5_ = 3.20, p = 0.024). A within-subjects comparison was used for the hM4Di data as few cells were successfully patched from control mice for this cohort. All data are presented as mean ± SEM. n = 4, 4 cells (hM3Dq); 6 cells (hM4Di).

**Extended Data Figure 9.**
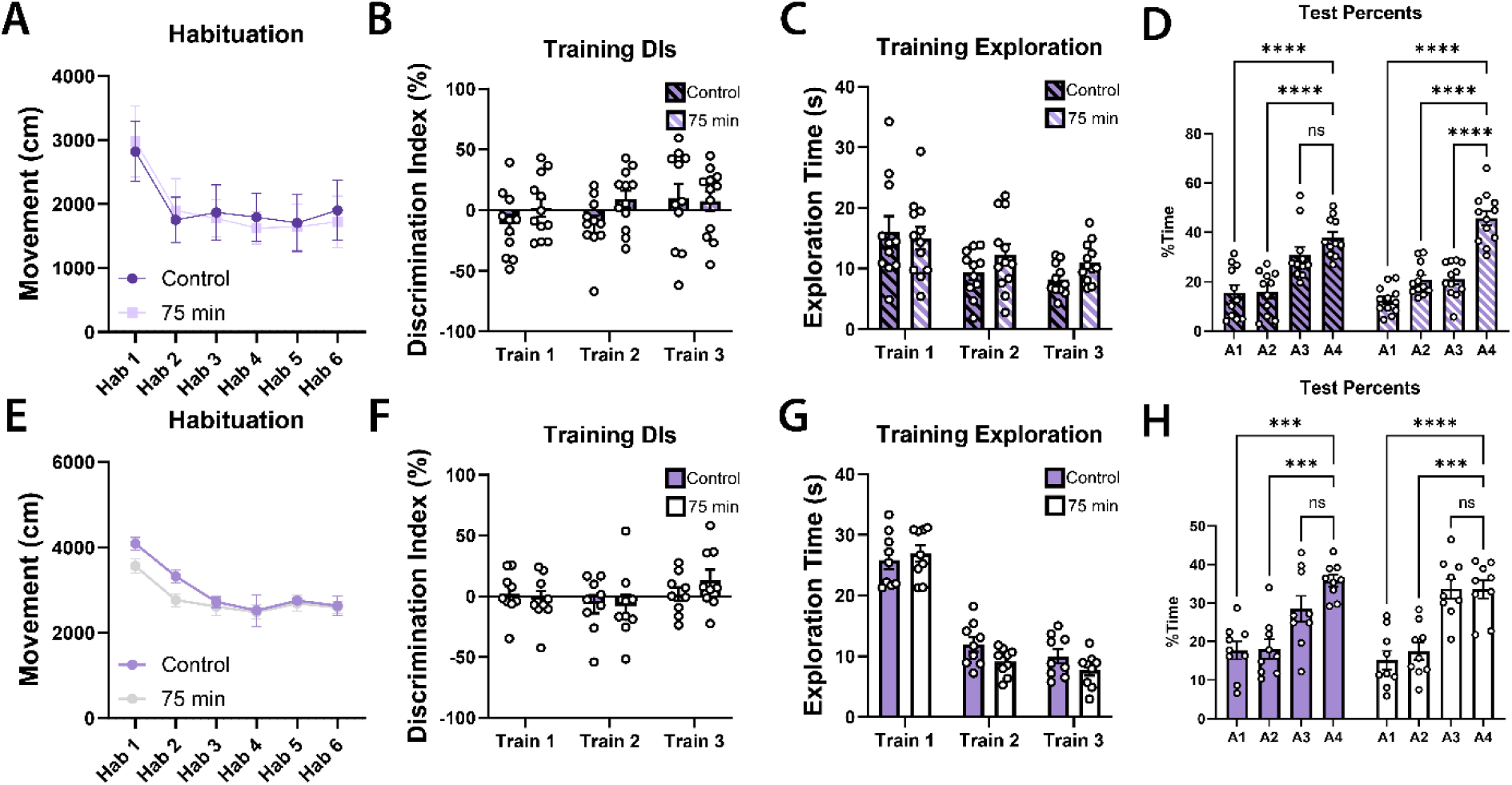
Additional data from the 75-min updating experiments (Figure 5). Old mice (**A**-**D**): **A.** Both groups habituated to the arenas as expected (two-way RM ANOVA: main effects of session and subject). **B.** Neither group had a significant object preference during the training sessions (two-way RM ANOVA, no significant effects). **C.** Both groups spent comparable time investigating objects during training (two-way RM ANOVA, effect of session). **D.** Investigation times at test displayed as percentages (two-way RM ANOVA, effect of object and object x group interaction; * denotes p < 0.05, ** denotes p < 0.01, *** denotes p < 0.001, **** denotes p < 0.0001 on Dunnett’s multiple comparisons test against A4). Young mice (**E**-**H**): **E.** Both groups habituated to the arenas as expected (two-way RM ANOVA: main effects of session and subject). **F.** Neither group had a significant object preference during the training sessions (two-way RM ANOVA, effect of subject). **G.** Both groups spent comparable time investigating objects during training (two-way RM ANOVA, effect of session). **H.** Investigation times at test displayed as percentages (two-way RM ANOVA, effect of object; * denotes p < 0.05, ** denotes p < 0.01, *** denotes p < 0.001, **** denotes p < 0.0001 on Dunnett’s multiple comparisons test against A4). All data are presented as mean ± SEM.

**Extended Data Figure 10.**
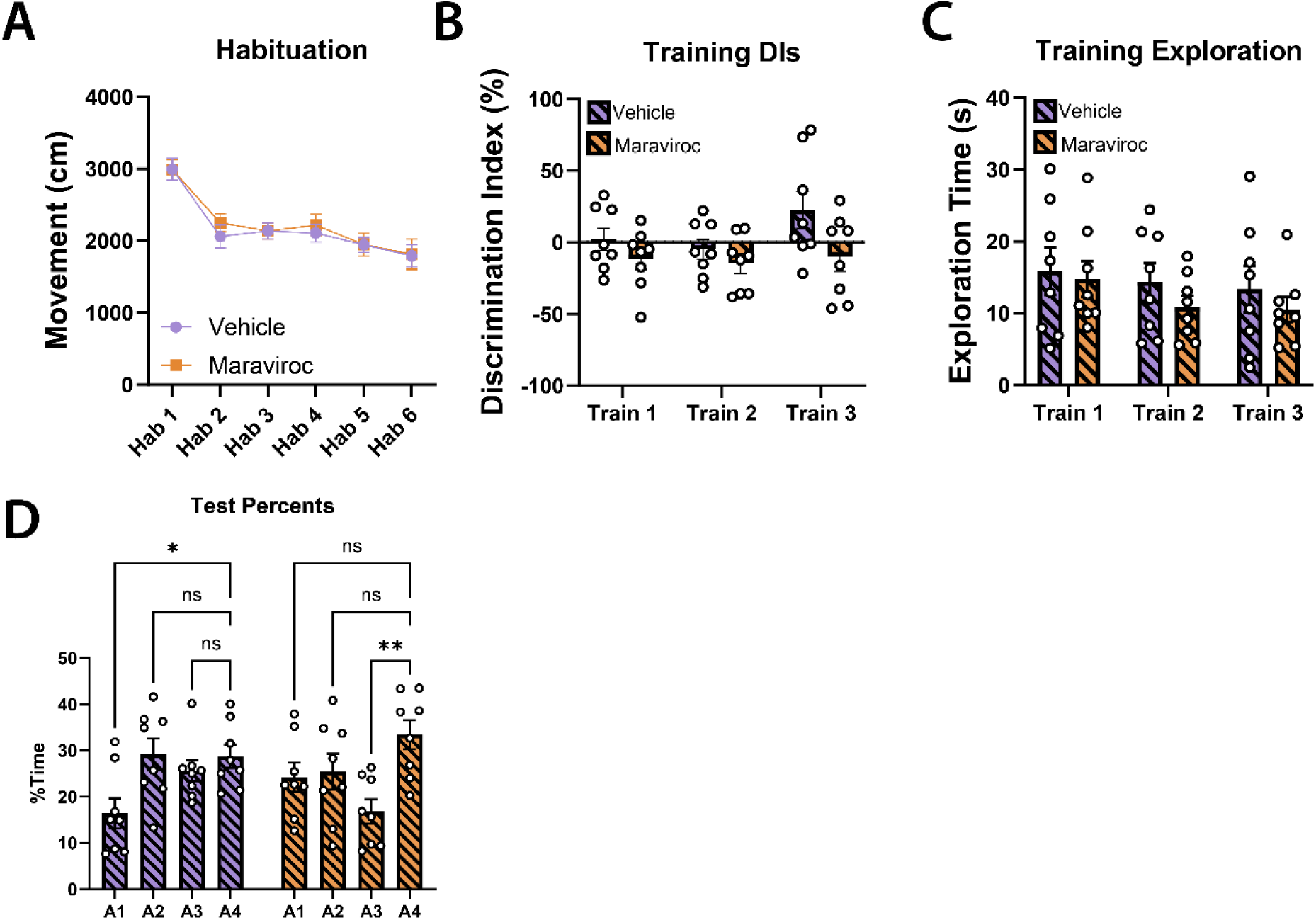
Additional data from the maraviroc experiment (Figure 6). **A.** Both groups habituated to the arenas as expected (two-way RM ANOVA: effects of session and subject). **B.** Neither group had a significant object preference during the training sessions (two-way RM ANOVA, effect of session). **C.** Both groups spent comparable time investigating objects during training (two-way RM ANOVA, effect of subject). **D.** Investigation times at test displayed as percentages (two-way RM ANOVA, effect of object; * denotes p < 0.05, ** denotes p < 0.01, *** denotes p < 0.001, **** denotes p < 0.0001 on Dunnett’s multiple comparisons test against A4). All data are presented as mean ± SEM.

## References

1. Wahlheim CN, Zacks JM. Memory updating and the structure of event representations. Trends Cogn Sci. 2025;29(4):380–392. doi:10.1016/j.tics.2024.11.008

2. Brunswick CA, Carpenter CM, Dennis NA, Kwapis JL. Not the same as it ever was: A review of memory modification, updating, and distortion in humans and rodents. Neurosci Biobehav Rev. 2025;174:106195. doi:10.1016/j.neubiorev.2025.106195

3. Przybyslawski J, Sara SJ. Reconsolidation of memory after its reactivation. Behav Brain Res. 1997;84(1):241–246. doi:10.1016/S0166-4328(96)00153-2

4. Nader K, Schafe GE, Le Doux JE. Fear memories require protein synthesis in the amygdala for reconsolidation after retrieval. Nature. 2000;406(6797):6797. doi:10.1038/35021052

5. Lee JLC, Nader K, Schiller D. An Update on Memory Reconsolidation Updating. Trends Cogn Sci. 2017;21(7):531–545. doi:10.1016/j.tics.2017.04.006

6. Jardine KH, Huff AE, Wideman CE, McGraw SD, Winters BD. The evidence for and against reactivation-induced memory updating in humans and nonhuman animals. Neurosci Biobehav Rev. 2022;136:104598. doi:10.1016/j.neubiorev.2022.104598

7. Rhodes MG. Age-Related Differences in Performance on the Wisconsin Card Sorting Test: A Meta-Analytic Review. Psychol Aging. 2004;19(3):482–494. doi:10.1037/0882-7974.19.3.482

8. Schoenfeld R, Foreman N, Leplow B. Ageing and spatial reversal learning in humans: Findings from a virtual water maze. Behav Brain Res. 2014;270:47–55. doi:10.1016/j.bbr.2014.04.036

9. St. Jacques PL, Olm C, Schacter DL. Neural mechanisms of reactivation-induced updating that enhance and distort memory. Proc Natl Acad Sci. 2013;110(49):19671–19678. doi:10.1073/pnas.1319630110

10. Weiler JA, Bellebaum C, Daum I. Aging affects acquisition and reversal of reward-based associative learning. Learn Mem. 2008;15(4):190–197. doi:10.1101/lm.890408

11. Kwapis JL, Alaghband Y, Keiser AA, et al. Aging mice show impaired memory updating in the novel OUL updating paradigm. Neuropsychopharmacology. 2019;45(2):337–346. doi:10.1038/s41386-019-0438-0

12. Mau W, Baggetta AM, Dong Z, et al. Ensemble remodeling supports memory-updating. bioRxiv. Preprint posted online March 23, 2023:2022.06.02.494530. doi:10.1101/2022.06.02.494530

13. Amelchenko EM, Bezriadnov DV, Chekhov OA, Anokhin KV, Lazutkin AA, Enikolopov G. Age-related decline in cognitive flexibility is associated with the levels of hippocampal neurogenesis. Front Neurosci. 2023;17. doi:10.3389/fnins.2023.1232670

14. Jardine KH, Minard EP, Wideman CE, et al. M1 muscarinic receptor activation reverses age-related memory updating impairment in mice. Neurobiol Aging. 2025;145:65–75. doi:10.1016/j.neurobiolaging.2024.10.007

15. Josselyn SA, Tonegawa S. Memory engrams: Recalling the past and imagining the future. Science. 2020;367(6473):eaaw4325. doi:10.1126/science.aaw4325

16. Reijmers LG, Perkins BL, Matsuo N, Mayford M. Localization of a stable neural correlate of associative memory. Science. 2007;317(5842):1230–1233. doi:10.1126/science.1143839

17. Zhou Y, Won J, Karlsson MG, et al. CREB regulates excitability and the allocation of memory to subsets of neurons in the amygdala. Nat Neurosci. 2009;12(11):1438–1443. doi:10.1038/nn.2405

18. Josselyn SA, Köhler S, Frankland PW. Finding the engram. Nat Rev Neurosci. 2015;16(9):9. doi:10.1038/nrn4000

19. Han JH, Kushner SA, Yiu AP, et al. Selective erasure of a fear memory. Science. 2009;323(5920):1492–1496. doi:10.1126/science.1164139

20. Liu X, Ramirez S, Pang PT, et al. Optogenetic stimulation of a hippocampal engram activates fear memory recall. Nature. 2012;484(7394):7394. doi:10.1038/nature11028

21. Roy DS, Arons A, Mitchell TI, Pignatelli M, Ryan TJ, Tonegawa S. Memory retrieval by activating engram cells in mouse models of early Alzheimer’s disease. Nature. 2016;531(7595):7595. doi:10.1038/nature17172

22. Park S, Kramer EE, Mercaldo V, et al. Neuronal Allocation to a Hippocampal Engram. Neuropsychopharmacology. 2016;41(13):13. doi:10.1038/npp.2016.73

23. Ramirez S, Liu X, Lin PA, et al. Creating a False Memory in the Hippocampus. Science. 2013;341(6144):387–391. doi:10.1126/science.1239073

24. Cai DJ, Aharoni D, Shuman T, et al. A shared neural ensemble links distinct contextual memories encoded close in time. Nature. 2016;534(7605):115–118. doi:10.1038/nature17955

25. Rashid AJ, Yan C, Mercaldo V, et al. Competition between engrams influences fear memory formation and recall. Science. 2016;353(6297):383–387. doi:10.1126/science.aaf0594

26. Shen Y, Zhou M, Cai D, et al. CCR5 closes the temporal window for memory linking. Nature. 2022;606(7912):7912. doi:10.1038/s41586-022-04783-1

27. Sehgal M, Filho DA, Kastellakis G, et al. Compartmentalized dendritic plasticity in the mouse retrosplenial cortex links contextual memories formed close in time. Nat Neurosci. 2025;28(3):602–615. doi:10.1038/s41593-025-01876-8

28. Mau W, Hasselmo ME, Cai DJ. The brain in motion: How ensemble fluidity drives memory-updating and flexibility. Colgin LL, ed. eLife. 2020;9:e63550. doi:10.7554/eLife.63550

29. Wright DS, Bodinayake KK, Kwapis JL. Investigating memory updating in mice using the Objects in Updated Locations (OUL) task. Curr Protoc Neurosci. 2020;91(1):e87. doi:10.1002/cpns.87

30. Ramirez S, Liu X, MacDonald CJ, et al. Activating positive memory engrams suppresses depression-like behaviour. Nature. 2015;522(7556):335–339. doi:10.1038/nature14514

31. Tsien JZ, Huerta PT, Tonegawa S. The Essential Role of Hippocampal CA1 NMDA Receptor–Dependent Synaptic Plasticity in Spatial Memory. Cell. 1996;87(7):1327–1338. doi:10.1016/S0092-8674(00)81827-9

32. Smies CW, Bellfy L, Wright DS, et al. Pharmacological HDAC3 inhibition alters memory updating in young and old male mice. Front Mol Neurosci. 2024;17. doi:10.3389/fnmol.2024.1429880

33. Gulmez Karaca K, Kupke J, Brito DVC, et al. Neuronal ensemble-specific DNA methylation strengthens engram stability. Nat Commun. 2020;11(1):639. doi:10.1038/s41467-020-14498-4

34. Gulmez Karaca K, Brito DVC, Kupke J, Zeuch B, Oliveira AMM. Engram reactivation during memory retrieval predicts long-term memory performance in aged mice. Neurobiol Aging. 2021;101:256–261. doi:10.1016/j.neurobiolaging.2021.01.019

35. Yiu AP, Mercaldo V, Yan C, et al. Neurons Are Recruited to a Memory Trace Based on Relative Neuronal Excitability Immediately before Training. Neuron. 2014;83(3):722–735. doi:10.1016/j.neuron.2014.07.017

36. Armbruster BN, Li X, Pausch MH, Herlitze S, Roth BL. Evolving the lock to fit the key to create a family of G protein-coupled receptors potently activated by an inert ligand. Proc Natl Acad Sci U S A. 2007;104(12):5163–5168. doi:10.1073/pnas.0700293104

37. Lau JMH, Rashid AJ, Jacob AD, Frankland PW, Schacter DL, Josselyn SA. The role of neuronal excitability, allocation to an engram and memory linking in the behavioral generation of a false memory in mice. Neurobiol Learn Mem. 2020;174:107284. doi:10.1016/j.nlm.2020.107284

38. Manvich DF, Webster KA, Foster SL, et al. The DREADD agonist clozapine N-oxide (CNO) is reverse-metabolized to clozapine and produces clozapine-like interoceptive stimulus effects in rats and mice. Sci Rep. 2018;8(1):3840. doi:10.1038/s41598-018-22116-z

39. Matos MR, Visser E, Kramvis I, et al. Memory strength gates the involvement of a CREB-dependent cortical fear engram in remote memory. Nat Commun. 2019;10(1):2315. doi:10.1038/s41467-019-10266-1

40. Cai DJ, Aharoni D, Shuman T, et al. A shared neural ensemble links distinct contextual memories encoded close in time. Nature. 2016;534 (7605):115–118. doi:10.1038/nature17955

41. Wilkin TJ, Gulick RM. CCR5 Antagonism in HIV Infection: Current Concepts and Future Opportunities. Annu Rev Med. 2012;63:81–93. doi:10.1146/annurev-med-052010-145454

42. Wang Y, Wu B, Jacob AD, et al. Neuronal competition shapes the encoding, consolidation, and retrieval of precise spatial memories in mice. Curr Biol. 2026;36(9):2255–2269.e6. doi:10.1016/j.cub.2026.03.058

43. de Sousa AF, Zeidler ZE, Almeida-Filho DG, et al. The prefrontal cortex controls memory organization in the hippocampus. Nat Neurosci. Published online April 28, 2026:1–12. doi:10.1038/s41593-026-02231-1

44. Gilboa A, Marlatte H. Neurobiology of Schemas and Schema-Mediated Memory. Trends Cogn Sci. 2017;21(8):618–631. doi:10.1016/j.tics.2017.04.013

45. Dunsmoor JE, Kroes MCW, Li J, Daw ND, Simpson HB, Phelps EA. Role of Human Ventromedial Prefrontal Cortex in Learning and Recall of Enhanced Extinction. J Neurosci. 2019;39(17):3264–3276. doi:10.1523/JNEUROSCI.2713-18.2019

46. St. Jacques PL, Olm C, Schacter DL. Neural mechanisms of reactivation-induced updating that enhance and distort memory. Proc Natl Acad Sci. 2013;110(49):19671–19678. doi:10.1073/pnas.1319630110

47. Schiller D, Kanen JW, LeDoux JE, Monfils MH, Phelps EA. Extinction during reconsolidation of threat memory diminishes prefrontal cortex involvement. Proc Natl Acad Sci. 2013;110(50):20040–20045. doi:10.1073/pnas.1320322110

48. Chowdhury A, Luchetti A, Fernandes G, et al. A locus coeruleus-dorsal CA1 dopaminergic circuit modulates memory linking. Neuron. 2022;110(20):3374–3388.e8. doi:10.1016/j.neuron.2022.08.001

49. Gálvez-Márquez DK, Salgado-Ménez M, Moreno-Castilla P, et al. Spatial contextual recognition memory updating is modulated by dopamine release in the dorsal hippocampus from the locus coeruleus. Proc Natl Acad Sci. 2022;119(49):e2208254119. doi:10.1073/pnas.2208254119

50. Wilson IA, Gallagher M, Eichenbaum H, Tanila H. Neurocognitive aging: prior memories hinder new hippocampal encoding. Trends Neurosci. 2006;29(12):662–670. doi:10.1016/j.tins.2006.10.002

51. Leal SL, Yassa MA. Neurocognitive Aging and the Hippocampus across Species. Trends Neurosci. 2015;38(12):800–812. doi:10.1016/j.tins.2015.10.003

52. Lee SH, Choi JH, Lee N, et al. Synaptic Protein Degradation Underlies Destabilization of Retrieved Fear Memory. Science. 2008;319(5867):1253–1256. doi:10.1126/science.1150541

53. Clem RL, Huganir RL. Calcium-Permeable AMPA Receptor Dynamics Mediate Fear Memory Erasure. Science. 2010;330(6007):1108–1112. doi:10.1126/science.1195298

54. Zhou M, Greenhill S, Huang S, et al. CCR5 is a suppressor for cortical plasticity and hippocampal learning and memory. eLife. 2016;5:e20985. doi:10.7554/eLife.20985

55. Bohnslav JP, Wimalasena NK, Clausing KJ, et al. DeepEthogram, a machine learning pipeline for supervised behavior classification from raw pixels. Mathis MW, Behrens TE, Mathis MW, Bohacek J, eds. eLife. 2021;10:e63377. doi:10.7554/eLife.63377

56. Vogel Ciernia A, Wood MA. Examining Object Location and Object Recognition Memory in Mice. Curr Protoc Neurosci. 2014;69(1). doi:10.1002/0471142301.ns0831s69

57. Kwapis JL, Alaghband Y, Kramár EA, et al. Epigenetic regulation of the circadian gene Per1 contributes to age-related changes in hippocampal memory. Nat Commun. 2018;9(1):3323. doi:10.1038/s41467-018-05868-0

58. López AJ, Kramár E, Matheos DP, et al. Promoter-Specific Effects of DREADD Modulation on Hippocampal Synaptic Plasticity and Memory Formation. J Neurosci. 2016;36(12):3588–3599. doi:10.1523/JNEUROSCI.3682-15.2016

59. Poll S, Mittag M, Musacchio F, et al. Memory trace interference impairs recall in a mouse model of Alzheimer’s disease. Nat Neurosci. 2020;23(8):952–958. doi:10.1038/s41593-020-0652-4

60. Joy MT, Ben Assayag E, Shabashov-Stone D, et al. CCR5 Is a Therapeutic Target for Recovery after Stroke and Traumatic Brain Injury. Cell. 2019;176(5):1143–1157.e13. doi:10.1016/j.cell.2019.01.044

61. Brocher J. biovoxxel/bv3dbox: BioVoxxel 3D Box - 1.24.7. Published online November 24, 2025. doi:10.5281/zenodo.17702242

62. Ting JT, Lee BR, Chong P, et al. Preparation of Acute Brain Slices Using an Optimized N-Methyl-D-glucamine Protective Recovery Method. J Vis Exp JoVE. 2018;(132):e53825. doi:10.3791/53825

63. Brunswick CA, Baldwin DJ, Bodinayake KK, et al. The clock gene Per1 is necessary in the retrosplenial cortex—but not in the suprachiasmatic nucleus—for incidental learning in young and aging male mice. Neurobiol Aging. 2023;126:77–90. doi:10.1016/j.neurobiolaging.2023.02.009

