## Supplemental Table 1 for "Old mice fail to integrate a memory update into an existing hippocampal engram"

Table S1. Primers and probes used for qPCR.

| **Oligo** | **Sequence** |
| --- | --- |
| *Ccr5* left primer | 5’-GTGCTGACATACCATAATCGATG-3’ |
| *Ccr5* probe | 5’-6-FAM/CCATCCTGC/ZEN/AAGAGCCAGAGTCTC /3IABkFQ-3’ |
| *Ccr5* right primer | 5’-TGTCTTCATGTTAGATTTGTACAGC-3’ |
| *Ccl5* left primer | 5’-CCTCTATCCTAGCTCATCTCCA-3’ |
| *Ccl5* probe | 5’-6-FAM/TCTTCTCTG/ZEN/GGTTGGCACACACTT  /3IABkFQ-3’ |
| *Ccl5* right primer | 5’-GCTCCAATCTTGCAGTCGT-3’ |
| *Gapdh* left primer | 5’-GGAGAAACCTGCCAAGTATGA-3’ |
| *Gapdh* probe | 5’-HEX/TCAAGAAGG/ZEN/TGGTGAAGCAGGCAT  /3IABkFQ-3’ |
| *Gapdh* right primer | 5’-TCCTCAGTGTAGCCCAAGA-3’ |
